# Automatically Defining Protein Words for Diverse Functional Predictions Based on Attention Analysis of a Protein Language Model

**DOI:** 10.1101/2025.01.20.633699

**Authors:** Hedi Chen, Jingrui Zhong, Xiaochun Zhang, Jingke Chen, Lin Guo, Xiaoliang Xiong, Xiaonan Zhang, Xiangyu Liu, Bailong Xiao, Boxue Tian

**Affiliations:** MOE Key Laboratory of Bioinformatics, State Key Laboratory of Molecular Oncology, Beijing Frontier Research Center for Biological Structure, School of Pharmaceutical Sciences, Tsinghua University, Beijing, 100084, China; Department of Natural Language Processing, Baidu International Technology (Shenzhen) Co., Ltd, Shenzhen 518000, China

**Keywords:** Protein function prediction, Protein words, Protein language model, Attention analysis, Community detection

## Abstract

Understanding the relationship between protein sequence and function remains a longstanding challenge in bioinformatics, and to date the lion’s share of related tools parse proteins at the domain or motif levels. Here, we define “protein words” as an alternative to “motif” for studying proteins and functional prediction applications. We first developed an unsupervised tool we term Protein Wordwise, which parses analyte protein sequences into protein words by analyzing attention matrices from a protein language model (PLM) through a community detection algorithm. We then developed a supervised sequence-function prediction model called Word2Function, for mapping protein words to GO terms through feature importance analysis. We compared the prediction performance of our protein word-based toolkit with a motif-based method (PROSITE) for multiple protein function datasets. We also assembled a functionally diverse data resource we term PWNet to support evaluation of protein words for predicting functional residues across 10 tasks (*e.g.*, diverse biomolecular binding, catalysis, and ion-channel activity). Our toolkit outperforms PROSITE in all the examined datasets and tasks. By abandoning domains and instead using attention matrices from a PLM for automatic, systematic, and annotation-agnostic parsing of proteins, our toolkit both outperforms currently available tools for functional annotations at the residue and whole-protein levels and suggests innovative forms of protein analysis well-suited to the post-AlphaFold era of biochemistry.

## Introduction

Understanding the relationship between protein sequence and function is a long-standing challenge in bioinformatics^1^. Protein functions have been annotated in terms of structure domains (*e.g.*, InterPro^2^, Pfam^3^), with functional annotations at higher resolution occurring at the motif level and individual residue level (for example through mutagenesis)^4,5^. Deep mutational scanning experiments can systematically characterize the function of individual residues in terms of some assayed function^6^. At the motif level, groups of residues hat associated with a specific protein function have been identified through structural analysis and using multiple sequence alignments (MSA)^7^.

Bioinformatics databases provide the functional annotations for protein sequences^8–11^. Machine learning models can leverage sequences, structures and biophysical descriptors to predict functional residues of proteins^12,13^. For example, a machine learning model combining statistical and biophysical properties is used to predict functional sites^14^. PhiGnet^15^ uses statistics-informed graph networks to predict protein functions, which assess the significance of residues for specific functions by characterizing evolutionary signatures. SPIRED-Fitness^16^ can predict both the 3D structure of a protein and its functional fitness from single sequence^17^. Protein language models (PLMs), such as ESM^18,19^ and ProtBert^20^, are powerful tools for protein function prediction^21–24^. For example, “CLEAN”^25^ uses ESM-1b with contrastive learning to predict enzyme functions.

The term “motif” has been frequently used in describing protein functions, though it is not well defined. Although we know that billions of proteins exert a myriad of functions, the number of proteins with experimental annotations at the residue- and motif-level remains relatively small, owing to the enormous work required for experimental bio-functional studies. Alva *et al.* (2015) previously explored an approach for evolutionary analysis that used an extended metaphor of “vocabulary”^26^, and identified a vocabulary of ancient peptides containing 40 “protein words” (with sequence lengths ranging from 9 to 40) through structural clustering. We were interested in applying such thinking to modern PLM-based protein analysis, and in developing an automatic tool to parse proteins into potentially informative semantic units, *i.e.*, “protein words”. Protein words can be applied to unique analytical capacities, such as functional residue prediction, functional annotations, studying the motions of proteins, studying signal transduction among protein complexes, and *de novo* protein design.

Computational prediction of protein words is attractive but challenging^21^. MSA-based methods are limited to intra-family studies and have difficulty with cross-family analysis^27^. PROSITE collects a set of human-curated residue patterns as rules for function annotation. However, it is slowly updated and often provides incomplete coverage for functional residue prediction. Supervised functional residue prediction models like PhiGnet can predict which residue is functional, however, they cannot predict detailed biochemical functions. Task-specific models such as small-molecule binding prediction method like CLAPE-SMB^28^, can only predict one function with one model.

In this study, we formally (and automatically) systematize semantic units of proteins as “protein words” by analyzing the attention matrices of a PLM through a community detection algorithm. We first present the unsupervised tool Protein Wordwise, which parses analyte protein sequences into protein words, and demonstrate the use of two “dictionaries” as filters that prioritize functionally informative protein words. We demonstrate use of Protein Wordwise for identifying the functional residues contributing to various known protein functions. We assembled a functionally diverse data resource we term Protein Words Net (PWNet) to support evaluation of protein words for predicting functional residues. We also developed a supervised sequence-function prediction model called Word2Function, for functional annotation word prediction through feature importance analysis. We show that our protein word-based toolkit outperforms the motif-based method (PROSITE), which is based on an enormous amount of human curation work. Given its automatic, systematic, and annotation-agnostic nature, our toolkit should facilitate investigations of protein motion, protein signaling interactions, and interpretations of functional contributions from structural features. Our study also demonstrates the extraction of new bio-information from attention matrices of large language models (LLMs), thus illustrating a new avenue for learning the knowledge within an LLM that could also be applied to other biological LLMs for DNA, RNA, and transcriptomics.

## Results

### Defining protein words based on analysis of attention heads in a PLM

Building on the previously explored concept of “protein words” from evolutionary analysis^26^, we propose extension of this term as an alternative to “motif” for functional prediction applications. BERT-based PLMs have been shown to perform well in predicting protein functions (EC numbers, GO terms) based on attention mechanisms^22,29^. Given the demonstrated high accuracy of reported PLMs for the predicting the functions of whole proteins, we speculated that analyzing residue relationships within attention heads of the PLM could in theory predict functionally informative protein words. We explored this using ESM2^18^, a powerful PLM trained on protein sequences in UniProt. Briefly, ESM2 has 33 layers and 20 heads at each layer (comprising a total of 660 attention heads). By inputting an analyte protein sequence into ESM2, we obtain 660 attention matrices from the corresponding attention heads. Each attention matrix is an *L*×*L* matrix, where *L* is the length of the input sequence (Fig. S1).

In a given attention matrix, the input protein is parsed as a set of residue pairs, and each pair is given an “attention value”, indicating a potentially informative interaction between a residue pair as predicted by an attention head. We use attention matrices as the input for predicting “protein words” (Fig. 1a). We define a “protein word” as a set of 5–20 residues. Given that available tools like unigram^30^ and N-gram^31^ are capable of informatively parsing analyte proteins into very short protein segments, we selected 5 as the lower bound for our protein word approach. Regarding the upper bound, Mandal et al. proposed 20 residues as the smallest number from which a fully protein-like folding motif can be formed^32^. Note that a protein word can be either contiguous or discontiguous in a protein sequence, and we allow a maximum of two gaps in a protein word. A gap can contain any number of residues (we depict a gap with an underline in the text and figures).

**Figure 1.**
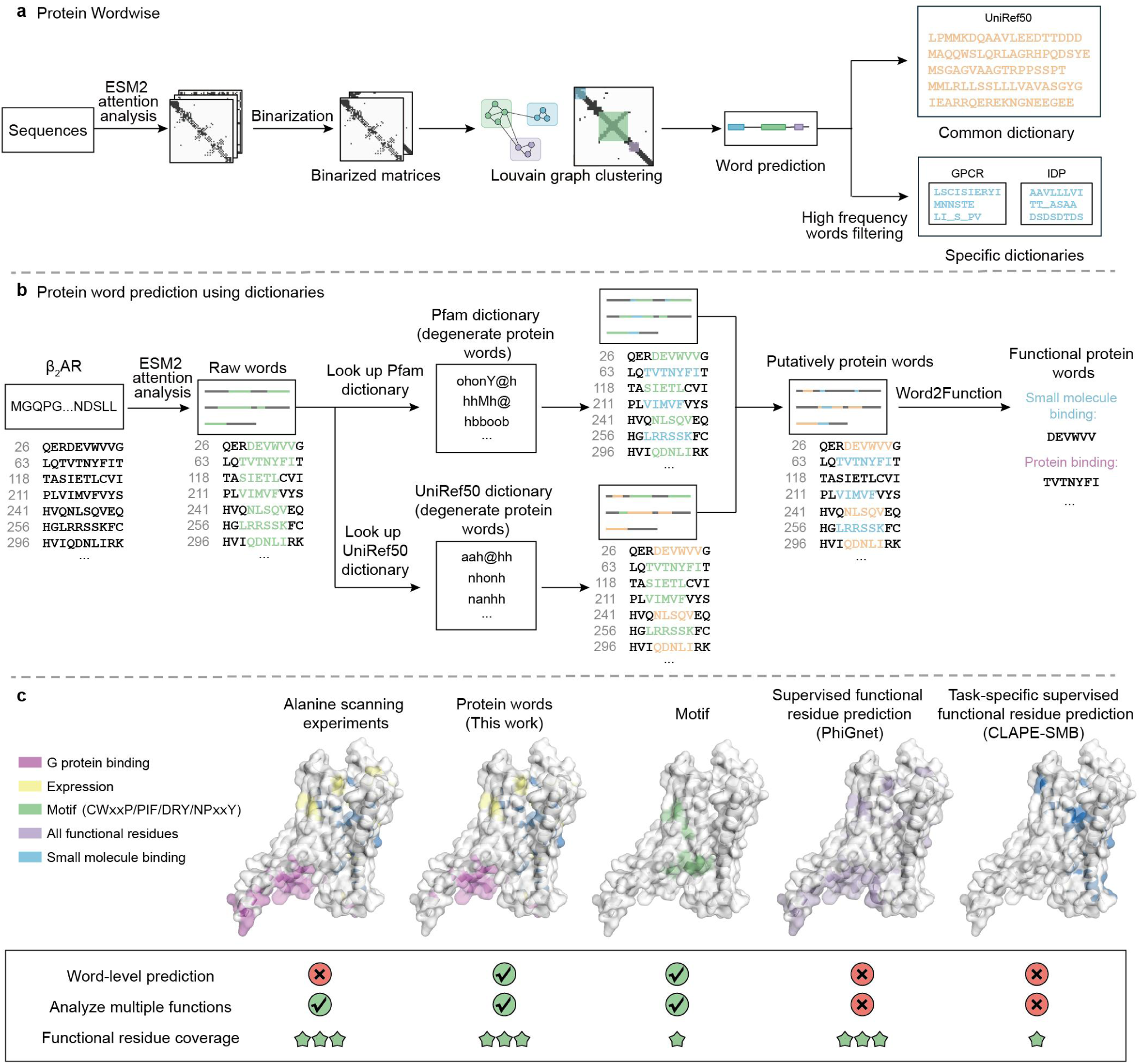
Prediction of protein words. **a**, The Protein Wordwise approach. We propose a protein word as an alternative to “motif” for functional prediction applications. A protein word is defined as a set of 5-20 residues in a protein, contiguous or with up to two gaps. The Protein Wordwise tool analyzes protein sequences using attention matrices. These matrices highlight potential residue interactions. We employ the Louvain community detection algorithm to identify protein words within these matrices. Upon analyzing multiple protein sequences, Protein Wordwise constructs dictionaries of high-frequency protein words. These dictionaries can then be used with a new analyte sequence to prioritize protein words likely to be functional. **b**, β_2_AR as an example analyte protein for dictionary-based word prediction. We constructed dictionaries by concatenating a set of commonly occurring words from multiple sequences with Protein Wordwise. Protein Wordwise takes a list of protein sequences as input, predicts raw words for each sequence, and identifies high-frequency words that occur in multiple sequences as the final output. We constructed two types of dictionaries: UniRef50 and a set of Pfam dictionaries. We converted the 20 residues into 12 degenerate residue types, and we convert raw words into their degenerate forms and then search for exact matches. **c**, Comparison of protein words with the other protein function annotation methods, using β_2_AR as an example analyte protein. Experimentally verified functional β_2_AR residues have been reported based on alanine scanning and structural analysis^35^. Protein Wordwise parses a sequence into protein words; residues within a protein word are putatively functional residues. Note that the potential function(s) of words are later assigned using our Word2Function tool.

### Predicting protein words with Protein Wordwise

We developed a protein word prediction approach called Protein Wordwise, which predicts protein words from an analyte sequence using a community detection algorithm (Fig. 1a and Methods). To aid analysis, note that we have compiled dictionaries of high-occurrence protein words initially detected in a large-scale analysis of UniProt sequences (Fig. 1b). These dictionaries are used to help prioritize high-occurrence protein words for the later step of functional annotation of protein words. Functional annotation task is performed by the “Word2Function” approach we developed. Notably, we use only primary sequences as input (*i.e.*, no structural information is employed). Upon combining Protein Wordwise and Word2Function, we achieve a word-level resolution for functional annotation of an analyte protein (Fig. 1b).

To predict protein words from attention matrices using Protein Wordwise, we binarize each attention matrix by defining two distinct cutoff values to determine which residue pairs are correlated vs. which residue pairs’ interactions can be considered as noise (Methods). This binary matrix can be represented as a graph, wherein nodes are residues and edges are residue-residue interactions^33^. We then use the Louvain algorithm^34^, a graph-based community detection algorithm, to segment the binary matrix into communities. Each community is considered as a “raw word”. The output for each attention matrix is a list of raw words, and we combine the output from all attention matrices, selecting raw words comprising between 5-20 amino acids (Fig. 1a). Note that one set of parameters is words lengths ranging from 5 to 6 residues in length while another parameter set is used for words of lengths ranging from 7 to 20 (Table S1).

The absence of published literature for protein words and dictionaries makes the task of assessing the accuracy of our approach for word-based functional annotation of proteins challenging. Not all residues within a protein sequence are functionally relevant. Among available public datasets for empirically examined functional residues, the DMS dataset comprises 198 protein sequences for which most residues were mutated to all of the other 19 possible residues (Fig. 2a). For the study that generated the DMS dataset, a residue was considered functional if at least one of these mutations influenced its function^11^; 37,033 out of 56,606 residues in the dataset were thusly defined as functional residues.

**Figure 2.**
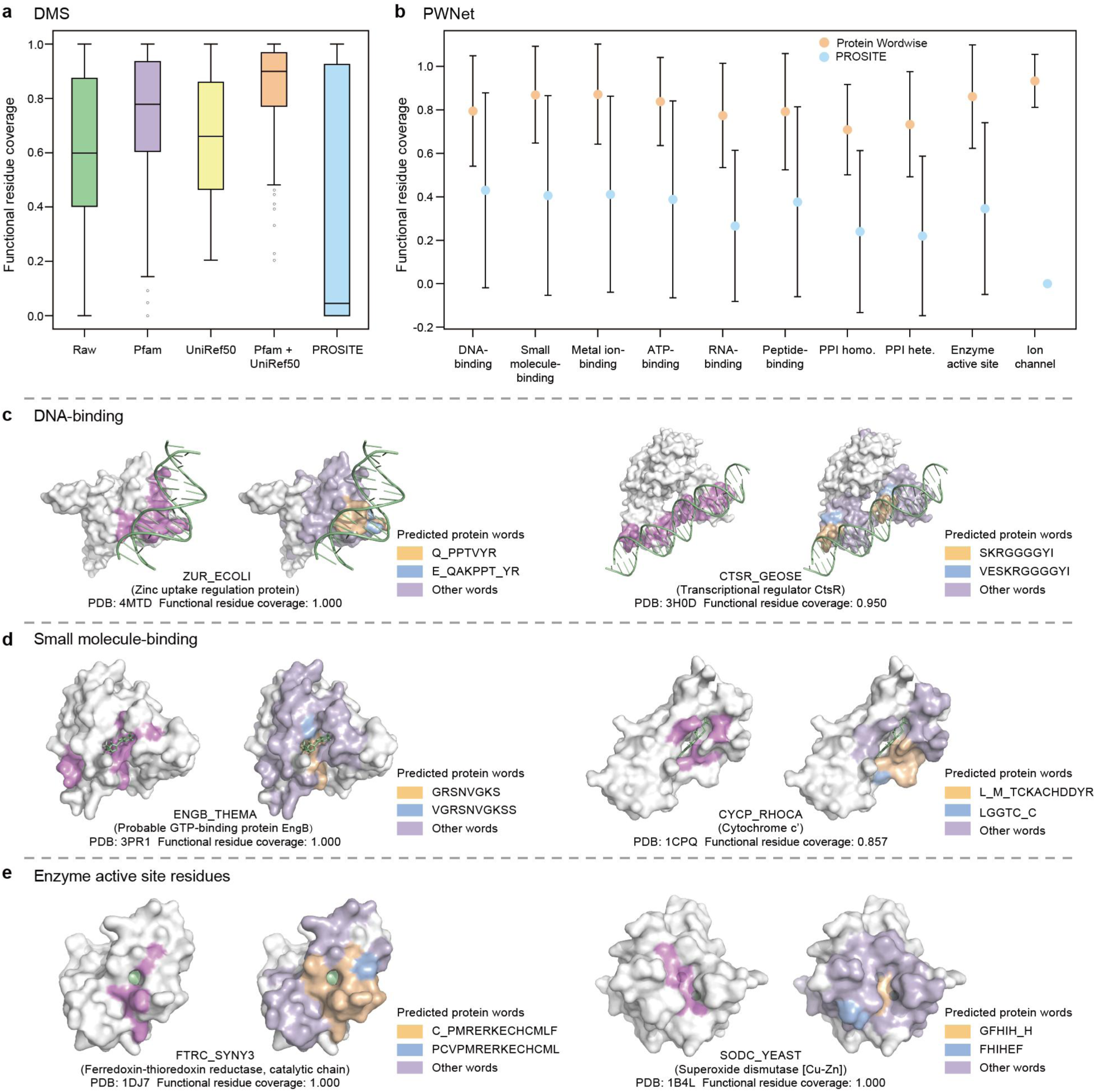
Protein words cover protein functional residues. **a,** Functional residue coverage for the DMS dataset using the indicated dictionaries. The median functional residue coverage for the raw words was 0.599. When incorporating Pfam, UniRef50, or both Pfam and UniRef50 dictionaries, the median functional residue coverage increased to 0.779, 0.661, and 0.900, respectively. **b,** Comparison of functional residue coverage for Protein Wordwise and PROSITE across 10 functional residue prediction tasks. We show that Protein Wordwise achieves higher functional residue coverage than the motif-based approach PROSITE for all the subsets in the PWNet dataset. **c**, Case studies on DNA-binding residue prediction. Left panel: the Zinc uptake regulation protein in *E. coli* (ZUR_ECOLI; PDB: 4MTD)^38^. 14 of ZUR’s 151 residues are known to function in DNA binding. The predicted protein words from Protein Wordwise covered all of ZUR’s binding site residues (functional residue coverage = 1.000). The functional residue coverage for PROSITE was 0.000 (cannot find a motif). Right panel: CtsR (CTSR_GEOSE; PDB: 3H0D); predicted protein words covered 19 of 20 known functional residues (functional residue coverage of 0.950; 0.000 for PROSITE) **d**, Case studies on small-molecule-binding residue prediction. Left panel: the GTP-binding protein EngB (ENGB_THEMA; PDB: 3PR1)^40^. 24 of 195 ENGB’s residues function in small molecule binding. The predicted protein words cover all binding residues. Right panel: cytochrome c’ (CYCP_RHOCA; PDB: 1CPQ). 7 of CYCP’s 129 residues function in small molecule binding. Protein Wordwise achieved a functional residue coverage of 0.857. **e**, Case studies on enzyme active site residue prediction. Left panel: the ferredoxin-thioredoxin reductase (FTRC_SYNY3; PDB: 1DJ7)^43^. 7 of FTRC’s 116 residues are enzyme active site residues. Protein Wordwise achieved a of functional residue coverage of 1.000. Right panel: the superoxide dismutase [Cu-Zn] (SODC_YEAST; PDB: 1B4L)^44^. 8 of SODC’s 154 residues are catalytic residues. All catalytic site residues are covered by predicted protein words (functional residue coverage = 1.000).

We can assess the prediction performance of Protein Wordwise based on “functional residue coverage”, which we define as the proportion of all functional residues covered by the predicted protein words for the analyte protein. To address false positives, we use “word accuracy”, which we define as the ratio of predicted words containing at least one functional residue to the total number of predicted words. As some false positive protein words may still be functionally relevant, we use functional residue coverage as our metric for Protein Wordwise.

### Using protein words and dictionaries for functional annotation of proteins

To illustrate use of protein words to investigate protein functions, we present β_2_AR as an example protein: this is a well-studied G protein-coupled receptor (GPCR) (Fig. 1c), for which functional residues have been experimentally verified based on alanine scanning^35^. Specifically, 195 out of 413 (47.2%) residues were experimentally determined to exert some function, such as adrenaline binding, G-protein binding, cholesterol binding, allosteric regulatory, and expression-level-related residues^35^. For such an analysis using Protein Wordwise and Word2Function, the query is the β_2_AR sequence, and the output is a set of protein words (Fig. 1b). Residues in final protein words are putatively functional residues. Potential function(s) of protein words are later assigned using Word2Function, *e.g.,* “small molecule binding” (Fig. 1b).

We initially predicted raw words for the β_2_AR receptor. Subsequently, we employed a GPCR Pfam dictionary and the large UniRef50 dictionary to “look up” protein words. This involves comparing each raw word to the dictionary entries to identify exact matches (Fig. 1b; Methods). In the β_2_AR example, Protein Wordwise predicted 186 out of 195 empirically defined functional residues (for a functional residue coverage of 0.954). Additionally, among the 208 words of this analyte protein, 184 words contained functional residues (for a word accuracy of 0.885). Given the automatic nature of Protein Wordwise, the semantic units (protein words) learned here can in theory be applied for studying protein-small molecule interactions, for protein motions, and for interpreting the functional contributions of structural features. Moreover, Protein Wordwise provides a new avenue for uncovering the hidden knowledge within an LLM.

### Degenerating words and compiling dictionaries

Given that Protein Wordwise outputs putatively functional protein words for each analyte protein, we assume that proteins with similar functions are likely to share functional protein words. If this is true, then it should be possible to use “dictionaries” of protein words as filters to enrich for functionally informative protein words. We constructed dictionaries by concatenating a set of commonly occurring words from multiple sequences with Protein Wordwise (Fig. S2b). It takes a list of protein sequences as input, predicts raw words for each sequence through attention analysis, and identifies high-frequency words that occur in multiple sequences as the final output (Fig. S2). We constructed two types of dictionaries: a common dictionary (derived from UniRef50^36^) and a set of family-specific dictionaries (predicted from each protein family defined by Pfam; currently, there are 20,762 family-specific dictionaries for Protein Wordwise). To collect words that can be used across different families and prevent biasing the Pfam with a large number of sequences (*e.g.,* PF02518 has 2 million sequences), the common dictionary is constructed from a collection of 1 million full length sequences from UniRef50: 50 randomly selected sequences representing each of the 20,000 Pfam families). Family-specific dictionaries (“Pfam dictionaries”) were constructed for each Pfam.

As our initial UniRef50 dictionary contains a large number of protein words (56,423,445), looking up words in such a dictionary is computationally intensive. Further, allowing for some aminoacidic diversity in the compiled dictionaries should support identification of relatively more functional protein words, as certain amino acids within a “word” can be substituted by residues having similar properties without altering the protein word’s function^37^. We degenerated the 20 proteinaceous residues into 12 residue types (*e.g.*, representing positively charged residues “R” and “K” as “b”; Table S2), which helped us reduce the UniRef50 dictionary size from 56,423,445 protein words to 2,581,748 degenerate protein words. Note that we also explored a more radical scheme in which the 20 proteinaceous residues were degenerated into only 4 residue types (Table S3). When looking up protein words in the dictionaries, we convert raw words into their degenerate forms and then search for exact matches (Fig. 1b).

We compared the functional residue coverage and word accuracy for dictionaries complied from the 20-, 12-, and 4-residue types, using the experimental data for β_2_AR, and we found that 12-residue type was the best choice in terms of the functional residue coverage and word accuracy values (Table S3, Table S4). Given this observed improved performance, we subsequently applied the 12-residue type scheme to all dictionaries (Table S1 for parameters). Before considering the potential emergent applications of this protein word approach, such as studying protein interactions, protein motions, interpreting the functional contributions of structural features, and protein design, we extensively investigated the use of protein words for tasks that current functional prediction tools can already achieve.

### Predicting functional residues of proteins with Protein Wordwise

For the functional residue prediction task, Protein Wordwise uses a single protein sequence as input (note we can also process each sequence of a batch [one-at-a-time]). We predict raw words and match raw words against the UniRef50 dictionary and a small number of suitable Pfam dictionaries (using annotated Pfams of the query sequences; Fig. 2a). When applied to 182 proteins in the DMS dataset with sequence lengths between 50 and 1,024 amino acids (removing peptides with lengths less than 50), the median and mean values for functional residue coverage using the combined Pfam+UniRef50 dictionaries were 0.900 and 0.843, respectively. We carried out ablation studies to assess dictionary contributions (Fig. S3). Upon removing the UniRef50 dictionary, the median and mean coverage decreased to 0.779 and 0.730, while removing all of the Pfam dictionaries resulted in a drop to 0.661 and 0.661. When relying solely on raw words for individual sequences (*i.e.*, without looking up in dictionaries), the median and mean coverage were obviously lower at 0.599 and 0.619.

To examine whether our approach outperforms PROSITE, a widely used motif-based function prediction approach, we performed a comparative analysis on functional residue coverage and word accuracy between Protein Wordwise and PROSITE. Of note, we found that PROSITE did not generate output for 35% of the protein sequences, likely because the number of rules are limited to a small number of sequences. The median and mean values for functional residue coverage using PROSITE were 0.045 and 0.393, which is much lower than those for Protein Wordwise (0.900 and 0.843). Protein Wordwise achieved 100% functional residue coverage for 16 of 182 examined proteins (*e.g.*, GAL4, GFP, see Table S5), while PROSITE achieved 100% functional residue coverage for 11 proteins. These findings support that Protein Wordwise is an effective tool for functional residue prediction.

### Predicting binding residues with Protein Wordwise

To test the use of Protein Wordwise for predicting functional residues in the context of molecular functions such as binding, we investigated DNA- and small-molecule binding tasks. We calculated the functional residue coverage for a dataset^28^ from a study that examined DNA-binding residues from 964 non-redundant protein sequences with experimental structures, for which 8.3% of the residues were classified as DNA-binding residues (Table S6). The median and mean functional residue coverage for protein words for this dataset when using Protein Wordwise were 0.921 and 0.795 (Fig. 2b). With PROSITE, the median and mean functional residue coverage values were 0.215 and 0.430.

We subsequently investigated two example DNA-binding proteins in greater detail (Fig. 2c). The first example is the Zinc uptake regulation protein in *E. coli* (ZUR_ECOLI), a transcriptional repressor known as highly sensitive to the Zn²^+^ concentration^38^. This protein consists of 151 residues, 14 of which function in DNA binding (22, 24, 26, 27, 44, 56, 58-60, 63, 64, 77, 82, 84). The predicted protein words from Protein Wordwise covered all of the known binding site residues for this protein (functional residue coverage = 1.000). The functional residue coverage for PROSITE was 0.000 (cannot find a motif). The second example protein is the transcriptional regulator CtsR in *G. stearothermophilus* (CTSR_GEOSE), which controls the expression of protein quality control genes by acting as a repressor of stress genes^39^. CTSR_GEOSE consists of 153 residues, 20 of which function in DNA binding (3-5, 27-29, 38, 40-41, 43-44, 49, 60-67). The predicted protein words cover 19 amino acids, with only residue 5 not covered, resulting in a functional residue coverage of 0.950. The functional residue coverage for PROSITE was 0.000. These results suggest that Protein Wordwise can effectively predict protein words for DNA-binding.

We then applied Protein Wordwise to a small-molecule-binding dataset^28^ comprising 4888 non-redundant full-length protein sequences, for which 3.3% of the residues were classified as small-molecule-binding residues (Table S6). The median and mean coverage values of protein words for all proteins of the dataset were 1.000 and 0.870 (Fig. 2b). We then studied the GTP-binding protein EngB in *T. maritima* (ENGB_THEMA), which is involved in ribosomes biogenesis^40^. ENGB_THEMA consists of 195 residues, 24 of which function in small molecule binding (30-37, 56-60, 74-77, 141-144, 173-175). The predicted protein words cover all binding site residues (functional residue coverage = 1.000) (Fig. 2d). The second example is cytochrome c’ in *R. capsulatus* (CYCP_RHOCA), a protein comprising 129 amino acids, 7 of which function in small molecule binding (10, 15, 68, 69, 118, 121, 122)^41^. The predicted protein words cover 6 of these amino acids (only residue 10 was not identified by Protein Wordwise), for a functional residue coverage of 0.857. These results show that Protein Wordwise is effective for predicting functional residues involved in DNA- & small molecule binding.

### Predicting functional residues across diverse tasks on PWNet

To test the use of protein words for predicting functional residues in the context of catalysis, we predicted enzyme active site residues for the 943 non-redundant proteins comprising the EasIFA dataset, wherein 1.5% of the residues were classified as active site residues^42^. Considering all proteins in this dataset, the mean functional residue coverage values for protein words was 0.861 (Fig. 2b), while the mean functional residue coverage for PROSITE was 0.346. We then specifically examined a ferredoxin-thioredoxin reductase in *Synechocystis sp.* (FTRC_SYNY3), which catalyzes the two-electron reduction of thioredoxins using electrons provided by reduced ferredoxin^43^. The protein consists of 116 amino acids, with 7 residues (56, 58, 75, 77, 86-88) constituting the enzyme active site. Protein Wordwise achieved a functional residue coverage of 1.000 for the active site residues. A second example is a superoxide dismutase [Cu-Zn] in *S. cerevisiae* (SODC_YEAST), which catalyzes the conversion of superoxide (O_2_^-^) into dioxygen and hydrogen peroxide^44^. This protein consists of 154 amino acids, with 8 residues (47, 49, 64, 72, 81, 84, 121, 144) serving as catalytic residues. All 8 catalytic site residues were covered by predicted protein words from Protein Wordwise (functional residue coverage = 1.000). The functional residue coverage for PROSITE was 0.375. These results support that protein words predicted by Protein Wordwise can be used to predict functional residues for catalysis.

We also generated a diverse dataset that we term PWNet to support evaluation of protein words for predicting functional residues across a range of tasks (Fig. 2b and Table S6). Very briefly, PWNet includes entries from publicly available datasets such as EasIFA and DIPS-Plus^45^ and represents 10 distinct functions including: identifying binding residues for protein-DNA (964 sequences), protein-RNA (915 sequences), protein-protein (8,238 sequences), protein-small molecule (4,888 sequences), protein-peptide (3,410 sequences), protein-ATP (390 sequences), and protein-metal ion interactions (3,485 sequences); residues involved in catalysis (943 sequences), and ion-channel activity (95 sequences) (Methods). PWNet entries have UniProt IDs and Pfam information, and an analysis of reported functional annotations indicates that the 10 functional subsets have average functional residue ratios ranging from 0.015 to 0.284 (Table S6). Protein Wordwise achieved a median functional residue coverage of 1.000 across 5 out of the 10 tasks. The mean functional residue coverage of these tasks ranged from 0.709 to 0.934, while the mean functional residue coverage for PROSITE ranged from 0.000 (ion channel) to 0.406. These results support that protein words can successfully predict residues performing a wide diversity of functions.

### Protein dictionary for predicting MHC peptides

MHC (Major Histocompatibility Complex) peptides-which are present on the surface of cells and are recognized by the immune system to identify foreign pathogens (*e.g.*, viruses, bacteria) and cancer cells^46^ (Fig. 3a)-are classified into two groups based on sequence length: MHC-I peptides (8 to 12 amino acids) and MHC-II peptides (13 to 25 amino acids)^47^. We conceptualize MHC peptides as naturally occurring protein words, and speculate that an ability to predict MHC-peptides could informatively guide the design of specific antigen epitopes to enhance vaccines efficacy. In current practice, predicting MHC peptides is often achieved using supervised learning models^48^. We expect that the UniRef50 dictionary contains many words that are identical to known MHC peptides, suggesting that we should be able to predict MHC-peptides in an unsupervised manner.

**Figure 3.**
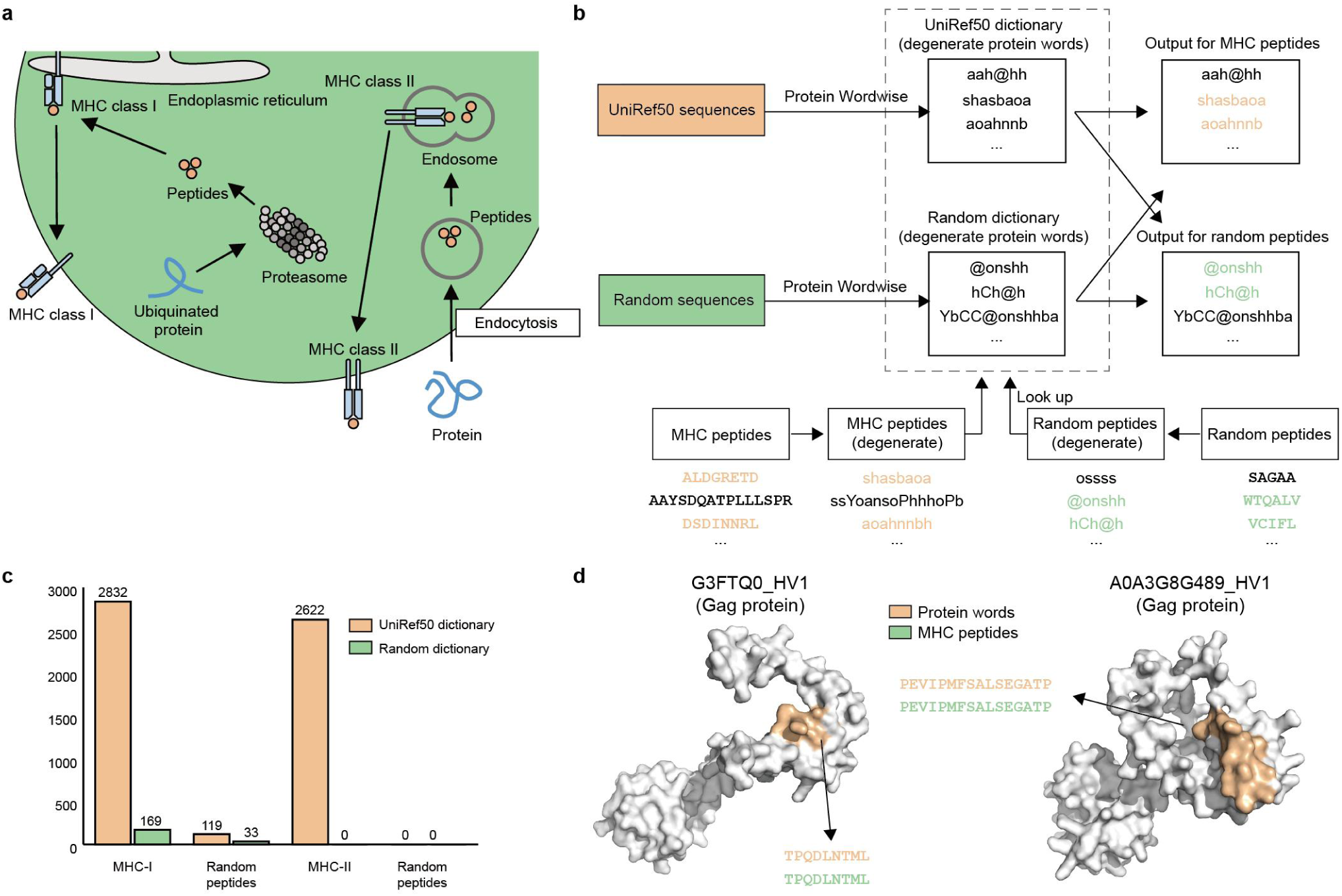
Protein dictionary for predicting MHC peptides. **a,** Schematic illustration of MHC peptide presentation. We conceptualize MHC peptides as naturally occurring protein words, with the UniRef50 dictionary expected to contain many words identical to known MHC peptides. **b,** Workflow for predicting MHC-peptides dictionaries. Both MHC-I peptides (8 to 12 amino acids) and MHC-II peptides (13 to 25 amino acids) were investigated. We converted MHC peptides into their degenerate forms and searched for exact matches with degenerate protein words from both the UniRef50 dictionary. As controls, a “random dictionary” was also generated from random protein sequences with the same sequence length distribution as UniRef50. We show that using the UniRef50 dictionary can find many more words than using the random dictionary. **c,** Comparison of UniRef50 and random words with MHC-I and -II peptides. Many UniRef50 predicted protein words are identical to MHC peptides (2,832 for MHC-I and 2,622 for MHC-II), while random words had virtually no overlap with MHC peptides. **d,** Case studies on two Gag proteins, with predicted protein words and known MHC-I peptides.

To explore the potential of using our UniRef50 dictionary for predicting MHC peptides, we converted MHC peptides into their degenerate forms and compared them with degenerate protein words from the UniRef50 dictionary. As control, a “random dictionary” was also compiled from randomly generated protein sequences of the same sequence length distribution as UniRef50 (Fig. 3b). We compare degenerate MHC-peptides against dictionaries to identify exact matches. The UniRef50 dictionary significantly outperformed the random dictionary for coverage of known MHC-I and MHC-II peptides (2,832 vs. 169 for MHC-I; 2,622 vs. 0 for MHC-II; Fig. 3c). These results indicate that the UniRef50 dictionary generated by Protein Wordwise can identify MHC peptides without requiring training data.

To gain insight into the sources of the matched protein words for MHC peptides in our Uniref50 dictionary, we extracted the set of Pfams of the original UniRef50 sequences that contributed protein words for any of the identified MHC peptides. Among the 1202 proteins containing MHC peptides and having experimental 3D structures, the top three Pfam domains were all Gag, with 48, 42, and 38 proteins, respectively (Fig. S4). Gag is a structural protein found in HIV-1 and other retroviruses. The MHC peptide “TPQDLNTML” was identified among the predicted protein words for G3FTQ0_HV1 in HIV-1. The MHC peptide “PEVIPMFSALSEGATP” was identified among the predicted protein words for A0A3G8G489_HV1 in HIV-1 (Fig. 3d). These successful predictions of MHC peptides from pathogens highlights Protein Wordwise’s potential for use in designing effective vaccines against infections.

### Mapping protein words to GO terms with Word2Function

Given that parsing proteins into words with Protein Wordwise supports the biological relevance of our functional residue prediction approach for 10 tasks on PWNet, we anticipated that the protein word approach would be able to provide informative annotations for protein functions at the word-level. We thus developed Word2Function, an ESM2-based supervised model that performs protein function prediction at the full-length protein level and that maps individual protein words to a curated subset of GO terms through feature importance analysis (Fig. 4a). Word2Function generates a GO-based “functional annotation word table” that can be leveraged for function prediction and potentially for protein design. We define a “functional annotation word” as a protein word with at least one function in our functional annotation word table.

**Figure 4.**
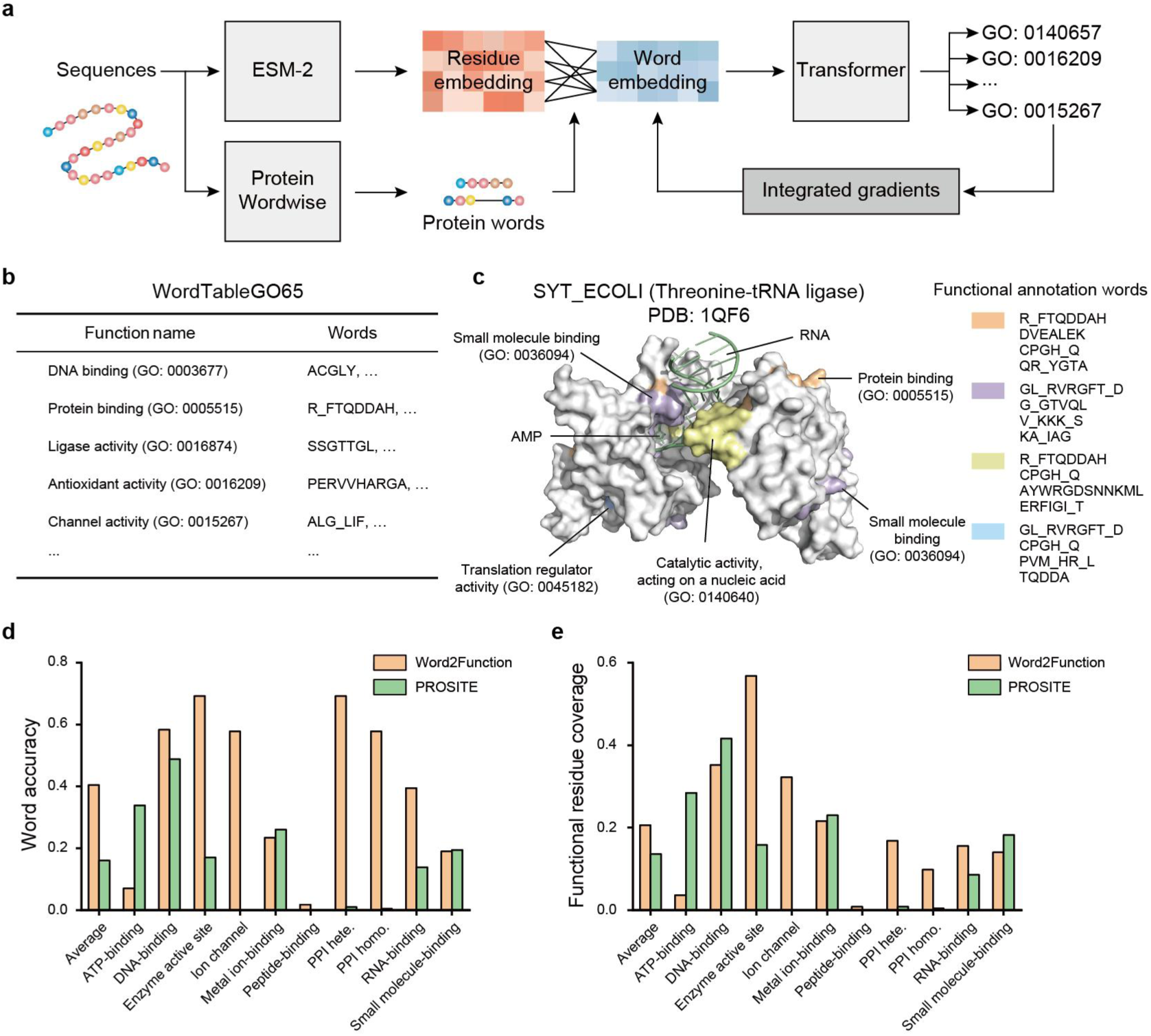
Mapping protein words to functions with Word2Function. **a**, Word2Function model overview. A residue embedding is a learned 1,280-dimensional vector representation of a residue within a protein sequence. Word2Function uses word embeddings, which are defined as the averaged embeddings of its constituent residue embeddings. The word embeddings are input into a transformer layer, followed by a linear layer to generate a probability vector for function prediction. During training, word embeddings are used to predict functions for the full-length protein sequences using ExpGO65 (GO terms). After training, an interpretability algorithm (Integrated Gradients) is employed to calculate the contribution of each word to different functions. Word2Function generates a “functional annotation word table” that can be leveraged for function prediction and potentially for protein design. Word2Function identifies the functional annotation words using a cumulative attribution score-based cutoff value α. The functional annotation words are then used to populate our functional annotation word table called “WordTableGO65”. **b**, The functional annotation word table (WordTableGO65) comprising 65 GO terms was generated by Word2Function. WordTableGO65 can be used for protein function prediction **c**, A case study of SYT_ECOLI. Threonine-tRNA ligase in *E. coli* (SYT_ECOLI) catalyzes the attachment of threonine to tRNA (Thr) and regulates translation^52^. Threonine-tRNA ligase associates with several GO terms, including “RNA-binding (GO: 0003723)”, “protein binding (GO: 0005515)”, “small molecule binding (GO: 0036094)”, and “translation regulator activity (GO: 0045182)”. Word2Function ranked a set of functional annotation words as relevant to these GO terms. The SYT_ECOLI example shows Word2Function’s ability to predict functional annotation words contributing to specific GO terms based on ranking of feature importance. Each GO term is associated with multiple functional annotation protein words. **d,e**, Comparison of word accuracy (**d**) and functional residue coverage (**e**) for Word2Function with PROSITE.

GO annotation is the most widely adopted method for protein function assignment^49^. The GO hierarchy organizes gene products, allowing for a systematic classification of their biochemical activities, with activities ranging from low-level categories like “ATP-dependent activity” to high-level categories such as “carbohydrate metabolism”. Our focus is on GO terms defined in Levels 1-3 in GO, that have experimental evidence. We constructed a dataset comprising 65 GO terms (Table S7; Methods), which we term ExpGO65. We first parsed all the 56,299 protein sequences in ExpGO65 into protein words by using Protein Wordwise, and then trained the ESM2-based Word2Function model to predict GO terms of the sequences.

ESM2-based models have achieved high accuracy in predicting GO terms using all residue embeddings of the protein sequence^50^, where a residue embedding is a learned 1,280-dimensional vector representation of a residue within a protein sequence. Instead of using residue embeddings, Word2Function uses word embeddings, which are defined as the averaged embeddings of its constituent residue embeddings (Methods). The word embeddings are input into a transformer layer, followed by a linear layer to generate a probability vector for function prediction (Fig. 4a). To link protein words to GO terms, we applied the Integrated Gradients algorithm^51^ to assess the influence of each protein word for the various GO terms (Methods).

Word2Function will determine the functional annotation words by using a cumulative attribution score-based cutoff value α. Word2Function ranks the word importance for each protein word based on its attribution score. Only protein words with a positive attribution score are ranked. A positive attribution score indicates that the input protein word positively contributed to the final prediction of GO terms. A cumulative attribution score is the sum of the positive attribution scores for all protein words in a sequence. We use a cutoff value, alpha, defined as the cumulative attribution score multiplied by a factor of 0.5. If the sum of attribution scores for the top-N protein words has attained this alpha value, we retain the top-N protein words as functional annotation words, discarding the other protein words. The functional annotation words are saved to our functional annotation word table called “WordTableGO65”.

### Illustrating the use of Word2Function for functional annotation word prediction

To illustrate the use of Word2Function for predicting functional annotation words, consider threonine-tRNA ligase in *E. coli* (SYT_ECOLI), which catalyzes the attachment of threonine to tRNA (Thr) and regulates translation^52^. Threonine-tRNA ligase associates with several GO terms, including “RNA-binding (GO: 0003723)”, “protein binding (GO: 0005515)”, “small molecule binding (GO: 0036094)”, and “translation regulator activity (GO: 0045182)”. Word2Function ranked a set of functional annotation words as relevant to “protein binding (GO: 0005515)”, including “R_FTQDDAH”, “DVEALEK”, “CPGH_Q” and “QR_YGTA” which an analysis of a solved SYT_ECOLI structure (PDB: 1QF6) showed as surface-exposed.

Protein words from SYT_ECOLI that were predicted as high importance for “small molecule binding (GO: 0036094)” included “GL_RVRGFT_D”, “G_GTVQL”, “V_KKK_S” and “KA_IAG”, all of which are positioned at a known AMP binding site or a surface pocket in the SYT_ECOLI structure. Protein words such as “R_FTQDDAH”, “CPGH_Q”, “AYWRGDSNNKML” and “ERFIGI_T” were ranked as high importance for “catalytic activity acting on a nucleic acid (GO: 0140640)” and these residues are positioned near the RNA binding site. Notably, the word “CPGH_Q” was observed in “protein binding (GO: 0005515)”, “translation regulator activity (GO: 0045182)” and “catalytic activity acting on a nucleic acid (GO: 0140640)”, demonstrating that Word2Function can assign multiple functions to a single functional annotation word. This SYT_ECOLI example shows Word2Function’s ability to predict functional annotation words contributing to specific GO terms based on ranking of feature importance.

### Comparing Word2Function with PROSITE

PROSITE uses manually curated rules based on motif-level annotation to predict protein functions, and is a widely used tool in bioinformatics applications^8^. To compare our Word2Function tool with PROSITE, we designed two tasks: Task1 is protein GO-term prediction at the whole-protein level; Task2 is prediction of functional annotation words/motifs. For Task1, we use mean functional Matthews Correlation Coefficient (MCC) as a metric, to address class imbalance in protein function prediction: a functional MCC is defined as the MCC for a binary classification problem to determine whether a protein sequence is associated with a specific GO term (Methods).

In Task1, across the 65 GO terms examined, Word2Function achieved a mean functional MCC of 0.424 on ExpGO65, with 28 GO terms exceeding a functional MCC of 0.500 (Table S8); while the mean functional MCC of PROSITE on ExpGO65 was 0.147. As Word2Function was trained using ExpGO65, testing on the same dataset may overestimate the model performance. We thus used the aforementioned PWNet as an external test set for comparing the performance of Word2Function and PROSITE for Task1. The mean functional MCC of Word2Function on PWNet was 0.284, while the mean functional MCC of PROSITE was 0.150.

For Task 2, we compared the mean functional residue coverage and mean word accuracy of Word2Function and PROSITE for PWNet. For Word2Function, the mean word accuracy was 0.202, and the mean functional residue coverage was 0.103; while for PROSITE, the mean word accuracy was 0.080, and the mean functional residue coverage was 0.068. Given that Word2Function outperforms PROSITE in both GO term prediction based on full-length sequence (Task1) and functional annotation word prediction (Task2).

### Case studies for functional annotation word prediction with Word2Function

We subsequently selected six proteins with diverse functions for case studies into how the protein word-level information provided by Word2Function can inform investigations into protein function, including: 3-hydroxydecanoyl-[acyl-carrier-protein] dehydratase in *E.coli* (FABA_ECOLI)^53^, type II methyltransferase M.*Pvu*II in *P. hauseri* (MTP2_PROHU)^54^, glutaredoxin 1 in *E.coli* (GLRX1_ECOLI)^55^, high mobility group protein HMG-I/HMG-Y in human (HMGA1_HUMAN)^56^, RNA-binding E3 ubiquitin-protein ligase in human (MEX3C_HUMAN)^57^ and vacuolar protein sorting-associated protein 4B in mouse (VPS4B_MOUSE)^58^. For these proteins, our predicted functional words achieved a mean word accuracy of 0.958 and a mean functional residue coverage of 0.661 (Fig. 5).

**Figure 5.**
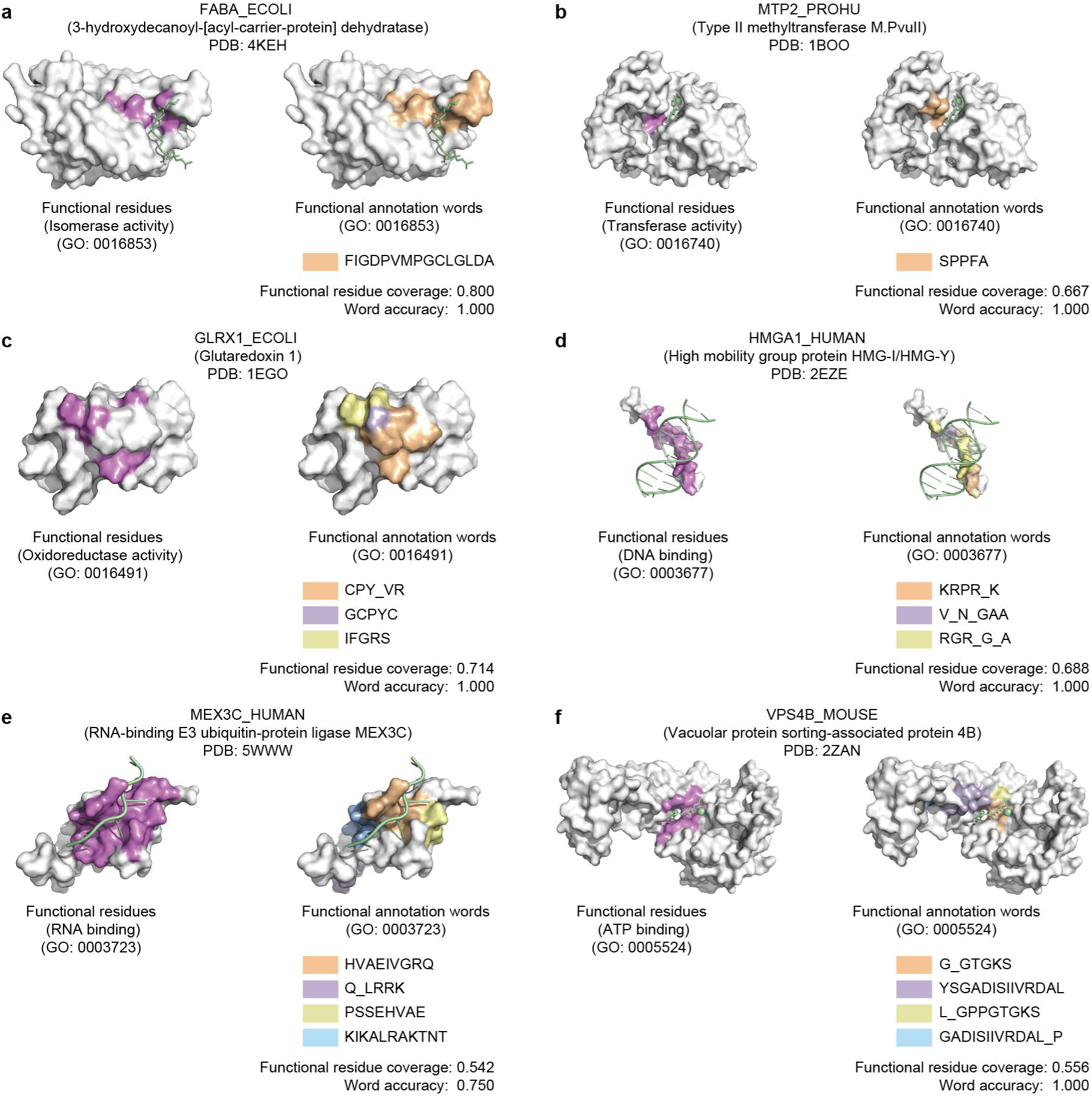
Predicting functional annotation words. We define a “functional annotation word” as a protein word with at least one assigned function in ExpGO65. We use six proteins with diverse functions as case studies to illustrate the use of Word2Function for identifying functional annotation words. The panels on the left display proteins highlighted with experimentally verified functional residues (violet); the panels on the right display proteins highlighted with predicted functional annotation words (orange and other colors). Word2Function predicts protein words that informatively map to specific functions and structural features of proteins. **a,** FABA_ECOLI (PDB: 4KEH)^53^. **b,** MTP2_PROHU (PDB: 1BOO)^54^. **c,** GLRX1_ECOLI (PDB: 1EGO)^55^. **d,** HMGA1_HUMAN (PDB: 2EZE)^56^. **e,** MEX3C_HUMAN (PDB: 5WWW)^57^. **f,** VPS4B_MOUSE (PDB: 2ZAN)^58^.

For FABA_ECOLI, an isomerase, Word2Function predicted two protein words related to “isomerase activity (GO: 0016853)”. The high importance word “FIGDPVMPGCLGLDA”, spans residues 72-86, among which residues V77, G80, and C81 were previously shown to form the catalytic triad (Fig. 5a). MTP2_PROHU, which methylates the 5’-CAGCTG-3’ sequence, had one predicted protein word (“SPPFA”) related to “transferase activity (GO: 0016740)”. This word spans residues 53-57, among which S53 and P54 are catalytic (Fig. 5b). These examples show that the functional word covers the known catalytic residues.

GLRX1_ECOLI is an oxidoreductase, and Word2Function predicted three protein words related to “oxidoreductase activity (GO: 0016491)”. The protein word “GCPYC” contains residues C11, P12, Y13, and C14, forming a disulfide bond between C11 and C14, that serves as an electron carrier (Fig. 5c). This example show that we can gain putative structural insights from Word2Function predicted words. HMGA1_HUMAN binds to DNA and is involved in the transcription regulation. Word2Function predicted three protein words related to “DNA binding (GO: 0003677)”, with word “KRPR_K” (spanning residues 55-58 and 65) ranked as the highest importance for this GO term. All residues in this word are functional residues and directly interact with DNA (Fig. 5d). MEX3C_HUMAN is involved in the post-transcriptional regulation of HLA-A allotypes. Word2Function predicted four protein words related to “RNA-binding (GO: 0003723)”, with word “HVAEIVGRQ” (spanning residues 31-39) ranked as the highest importance for this GO term. Structurally, residues V32, A33, E34, V36, G37, R38, and Q39 directly interact with RNA (Fig. 5e). VPS4B_MOUSE, which is involved in the endosomal MVB pathway, had four protein words related to “ATP-binding (GO: 0005524)”, with the “G_GTGKS” protein word comprising residues 51 and 54-58, among which residues G54, T55, G56, K57, and S58 interact with ATP (Fig. 5f). The examples of MEX3C_HUMAN and VPS4B_MOUSE show that the functional word covers the known RNA-binding and ATP-binding residues. These results show that Word2Function predicts protein words that informatively map to specific, known functions and structural features of analyte proteins.

### Using WordTableGO65 for full sequence-level protein function prediction

Given the WordTableGO65 generated using Word2Function provides protein word-level information, reusing these words as “rules” for function prediction is an attractive potential application. That is, we should be able to use Protein Wordwise to parse sequence into protein words and then use WordTableGO65 to find exact matches with these protein words. Given that WordTableGO65 was populated based on sequences/whatever of known function, thusly assigning a given protein word to a corresponding entry in WordTableGO65 would represent a word-based annotation of an analyte protein sequence. We evaluated our WordTableGO65 for Task1 on ExpGO65, and achieved an average functional MCC of 0.498 (Table S9), which is higher than the MCC of 0.424 for Word2Function. We also tested the performance of the WordTableGO65 for Task1 on PWNet. The functional MCC of WordTableGO65 on PWNet was 0.286, which is slightly higher than that of Word2Function (0.284). We subsequently used WordTableGO65 for additional whole-protein level function prediction applications.

We then compared the mean functional residue coverage and mean word accuracy of Word2Function and this WordTableGO65 approach for Task2 on PWNet. Word2Function achieved higher values for both metrics (mean word accuracy: 0.202, and the mean functional residue coverage: 0.103 (vs. 0.193 and 0.086 for WordTableGO65). Thus, WordTableGO65 can be used as an effective tool for functional annotation at the full-length sequence level, but does not perform as well as Word2Function for function prediction at the protein word-level.

### Predicting functions of DUFs with Protein Wordwise, Word2Function, and WordTableGO65

The term “domain of unknown function” (DUF) is used in biological data resources for a protein domain that has no characterized function^59^, and DUFs currently comprise ∼2% of the UniProt. It is notable that currently available supervised protein functional annotation models typically perform poorly for DUFs. We concatenated over one million sequences annotated as “DUF” in the InterPro database as a dataset we term “InterProDUF”. We first used Protein Wordwise to parse DUFs into protein words. We then searched for matches between these predicted protein words for DUFs and entries in our WordTableGO65 (Fig. 6a). GO terms related to DUFs were thusly predicted, and functional annotation words were identified. The most prevalent predicted functions for DUF sequences included protein binding, hydrolase, and transferase activities (Fig. 6b). The high resolution functional annotation obtained with our approach can in theory guide mutagenesis experiments to focus on potentially fruitful regions for investigating the functions of DUFs.

**Figure 6.**
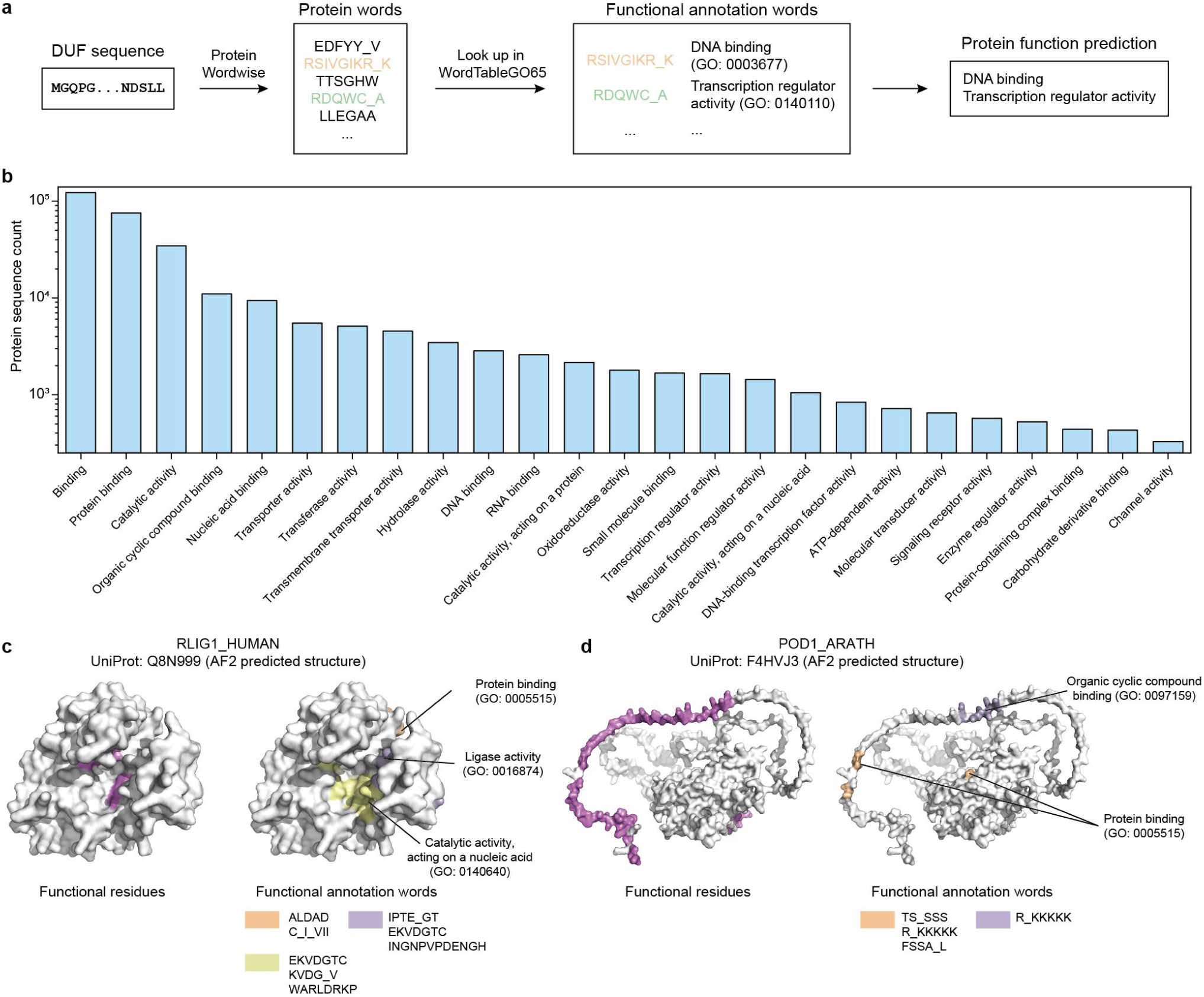
Predicting functions and functional annotation words for DUFs. **a,** Function prediction for DUFs by using WordTableGO65. **b,** Distribution of Top-25 predicted functions for DUFs. **c,** A case study on the DUF sequence RLIG1_HUMAN in human^60^. WordTableGO65 identifies 7 functional annotation words. The associated GO terms for these words suggest functions related to “ligase activity (GO: 0016874)”, “catalytic activity, acting on a nucleic acid (GO: 0140640)” and “protein binding (GO: 0005515)”. Site-directed mutagenesis of a recent experimental study support our prediction. **d,** A case study on DUF sequence POD1_ARATH in *A. thaliana*^61^. WordTableGO65 identified 3 functional annotation protein words within POD1_ARATH. The associated GO terms for these words suggested functions related to “protein binding (GO: 0005515)” and “organic cyclic compound binding (GO: 0097159)”. A recent experimental study showed that POD1_ARATH was involved in retaining proteins within the endoplasmic reticulum lumen. Further, mutation experiments indicate that the N-terminal 60 residues of POD1, enriched in Lys/Arg motifs, function in endoplasmic reticulum retention. Notably, the predicted word “R_KKKKK” was associated with both “protein binding” and “organic cyclic compound binding”.

As a case study, we examined the DUF sequence RLIG1_HUMAN in humans^60^. Protein Wordwise identified 105 “protein words” within RLIG1_HUMAN, of which 7 were found in WordTableGO65 (Fig. 6c). The associated GO terms for these words suggest functions related to “ligase activity (GO: 0016874)”, “catalytic activity, acting on a nucleic acid (GO: 0140640)”, and “protein binding (GO: 0005515)”. A recent experimental study indicated that RLIG1_HUMAN functions as an RNA ligase, and site-directed mutagenesis has shown that T55, K57, D59, and R77 are crucial for its RNA ligase activity. Within RLIG1_HUMAN, several “functional annotation words” are linked to the mutation sites. Specifically, “IPTE_GT” (53-56, 60-61) is associated with “ligase activity”, covering T55; “EKVDGTC” (56-62) is linked to both “ligase activity” and “catalytic activity, acting on a nucleic acid”, covering K57 and D59; “KVDG_V” (57-60, 65) is associated with “catalytic activity, acting on a nucleic acid”, covering K57 and D59; and “WARLDRKP” (75-82) is also linked to “catalytic activity, acting on a nucleic acid”, covering R77.

As another case study, we examined the DUF sequence POD1_ARATH in *A. thaliana*^61^ (Fig. 6d). WordTableGO65 identified 3 functional annotation words for POD1_ARATH. The associated GO terms for these words suggested functions related to “protein binding (GO: 0005515)” and “organic cyclic compound binding (GO: 0097159)”. A recent experimental study indicated that POD1_ARATH is likely a component of the calreticulin 3 (CRT3) complex, potentially functioning as a co-chaperone involved in retaining proteins within the endoplasmic reticulum lumen. Further, mutation experiments from that study revealed that the N-terminal 60 residues of POD1, enriched in Lys/Arg motifs, function in endoplasmic reticulum retention. Notably, the predicted word “R_KKKKK” (53-59) was associated with both “protein binding (GO: 0005515)” and “organic cyclic compound binding (GO: 0097159)”. These results expand beyond our earlier analyses of proteins of known function, demonstrating that the functional word table can predict functions for DUF proteins and uncover new functional sites. Finally, and more broadly, while our results establish that our protein word toolkit performs very well for functional prediction tasks that currently available protein functional prediction approach also support, we are most excited about the uniquely emergent opportunities from this automated and annotation-agnostic approach for parsing proteins into semantic units.

## Discussion

In this study, we defined protein words based on analysis of attention matrices of a PLM using a community detection algorithm. We developed Protein Wordwise for parsing protein sequences into protein words. We constructed UniRef50 and Pfam dictionaries to use as filters to prioritize functionally informative protein words. We assembled the PWNet dataset to support evaluation of protein words for predicting functional residues. We also developed Word2Function, which maps protein words to GO terms and populates a functional annotation word table called WordTableGO65. We demonstrate that our toolkit outperforms the motif-based method PROSITE in terms of function prediction using the PWNet dataset.

A large number of proteins in the UniProt database, despite possessing functions, lack experimental annotations at a higher resolution than “protein domains”. Residue- and motif-level annotation often requires extensive human labor (and expertise). For understanding and designing protein function, predicting functional protein words may enable a “functional group” concept analogous to how small-molecule functional groups are utilized in chemistry (*e.g.*, carboxyl groups are acidic). Ultimately, we aspire to build a knowledge base of functional protein words, and the functional annotation word table presented in this work constitutes a good starting point for this knowledge base. As the knowledge base of protein words expands, our understanding of informative and practically actionable protein characteristics will likely increase. Such knowledge should aid in the rational modulation of protein function by guiding the design of diverse sequence mutations (likely at the protein word level; *i.e.*, beyond single residue). We can identify shared words across a set of proteins with a specific feature, *e.g.,* proteins that can undergo liquid-liquid phase separation in cells. These shared words can then be harnessed for protein design tasks. Further work will be needed to assess the accuracy and generalizability of predicted functional protein words, including for applications in protein design (currently under investigation).

Representing proteins with protein words may also aid in understanding protein motion, which could support sophisticated studies of dynamic biological systems. For example, consider that for studying signal transduction interactions between (and within) protein complexes, residue-level analysis is often insufficient^62^. Using protein words as proposed here may enable a middle ground for investigating signaling interactions that lies between extremely complex atomic resolution models and relatively unsophisticated domain-level models, potentially yielding insightful predictions at a resolution that is well-suited to engineering and design applications. Moreover, it should be relatively straightforward to use functional protein words for constructing coarse-grained models to study protein dynamics. Current coarse-grained MD simulations primarily focus on the functional groups within an amino acid^63^. Our approach can automatically parse a sequence into protein words, which can facilitate the segmentation of the protein into functionally informative groups for building coarse-grained models for proteins that are tailored to specific functional questions. Such models would likely assist in the interpretation of functional contributions from structural features from multiple proteins, and use of relatively simpler models would, computationally, empower researchers by allowing for longer simulation time scales.

The Uniref50 and Pfam dictionaries, as well as WordTableGO65 can be viewed as a set of semantic rules for proteins. The spirit of our approach is that we learn these rules by analyzing the attention matrices of the ESM2 model, without any human curation (distinct from PROSITE). This approach provides a new avenue for learning the knowledge within a large language model. We therefore anticipate that our general approach should be applicable for other foundational biological large language models, such as DNA^64^, RNA^65^ and transcriptomics^66^, to discover the “words” for DNA, RNA, and for molecular genetics pathways.

## Methods

### Overview

A protein word is defined as a set of 5-20 residues in a protein sequence. A protein word can be either contiguous or discontiguous in a protein sequence. We developed an unsupervised protein word prediction tool we term Protein Wordwise, and we compiled the Uniref50^9^ dictionary and a set of Pfam^3^ dictionaries. These dictionaries are used to help prioritize high-occurrence protein words for the later step of functional annotation of protein words. The Uniref50 dictionary is also used to identify MHC peptides. We developed a functional word prediction tool we term “Word2Function”. We use Word2Function to map protein words to functions, and then populate a functional annotation word table from Word2Function output that we term “WordTableGO65” by training a supervised model based on the ExpGO65 Dataset. By combining Protein Wordwise, Word2Function, and the WordTableGO65, we can predict protein functions at the word level (Fig. 1). We compared the prediction performance of our toolkit against a motif-based approach (PROSITE)^8^ using the DMS^11^ and PWNet datasets. We show that our approach outperforms PROSITE in protein function prediction at both the full-sequence level and word-level. Finally, we show that the functional annotation word table (WordTableGO65) supports function predictions for DUFs.

### Protein Wordwise

#### Attention matrix preparation

Protein sequences were downloaded from UniProt (2024/01)^9^, and Pfam (version 36.0)^3^ was used for all Pfam-related tasks. ESM2^18^, a widely used PLM, was used throughout the study. As the ESM2 model only supports sequence length up to 1024, sequences longer than 1024 were discarded. Sequences with noncanonical or unknown residues were deleted from the dataset. The original ESM2-650M variant was run for all input sequences (without specific finetuning) to generate attention matrices (660 matrices for each sequence). Each matrix was converted to a binary matrix.

We first calculate the maximum and minimum values of the attention matrix, and then set the lower bound as 10% of the range of values of the attention matrix. The attention values lower than this lower bound were set to 0, while the other values were set to 1. If the number of 1 in the attention matrix exceeds 40% of the total number of all residue pairs, we will calculate the upper bound, which is defined as the value that accounts for 60% of all attention values. The attention values lower than the the upper bound were set to 0.

#### Raw word prediction via Louvain community detection algorithm

The binary matrix was converted to a directed graph, wherein nodes are the residues, and an edge from one node to another exists if and only if the binarized attention value between the two residues is 1. Based on such directed graph representation of attention matrices, we apply the Louvain community detection algorithm^34^ with resolution parameter γ=1.0 and modularity gain threshold ΔQ=10^-7^ here for residue nodes clustering. All the clusters were initially filtered by a length range of 5 to 20. Each residue cluster (community) we term a “raw word”. High occurrence raw words for a given set of sequences (*e.g.,* a Pfam family) were used to compile a dictionary.

#### Compiling Pfam and UniRef50 dictionaries (multi-sequence mode)

Two word prediction modes were developed: single-sequence mode and multiple-sequence mode. A multi-sequence mode must be run initially to generate dictionaries for subsequent use. Two types of dictionaries were constructed: a common dictionary (derived from UniRef50^36^) and a set of family-specific dictionaries (predicted from each protein family defined by Pfam^3^; currently, 20,762 Pfam dictionaries exist for Protein Wordwise). In the multi-sequence mode, Protein Wordwise accepts a list of protein sequences as input, and predicts raw words for each sequence. A common dictionary was then constructed by collecting raw words of 1 million full-length sequences from UniRef50: 50 randomly selected sequences representing each of the 20,000 Pfam families. Family-specific dictionaries (“Pfam dictionaries”) were constructed for each Pfam family. For each raw word, the 20 proteinaceous residues were degenerated into 12 residue types (*e.g.*, representing positively charged residues “R” and “K” as “b”; Table S2). We stratified the dictionary by length, retaining high occurrence raw words for each length. The percentage thresholds for length 5 to 20 are: 5% for 5-residue, 2.5% for 6-residue, 0.15% for 7 to 11-residue, and 0.3% for 12 to 20-residue. These parameters were obtained by evaluating the dictionary performance with the functional residue coverage and word accuracy on the β_2_AR (Table S1).

#### Prioritize high-occurrence protein words for functional annotation of protein words (single sequence mode)

With the dictionaries generated from the multi-sequence model, we use the single-sequence for predicting protein words. The single-sequence mode uses a single protein sequence as input, and we can also process each sequence of a list one-at-a-time). We first predict raw words using attention analysis described in previous steps. We then look up raw words in the UniRef50 and Pfam dictionaries (Fig. 2a). When looking up protein words in the dictionaries, we convert raw words into their degenerate forms and then search for exact matches (Fig. 1b). In a protein word prediction task for a single sequence, when the query protein sequence has a related Pfam dictionary, we use UniRef50+Pfam dictionaries by default (Fig. 1b). For sequences without a related Pfam dictionary, we use the UniRef50 dictionary to look up protein words.

#### MHC peptide prediction by using the UniRef50 dictionary

MHC peptides were obtained from the MHC Motif Atlas dataset^47^. 679,294 MHC I peptides and 1,173,480 MHC II peptides were converted into 325,384 and 404,696 unique 12-residue degenerate forms, respectively. For MHC I, we focused on sequences between 8 and 14 amino acids in length based on the length distribution for this joined dataset; while for MHC II, we considered sequences between 12 and 20 amino acids. To investigate whether protein words are better than random sequences in terms of MHC peptide prediction, we generated an equal number of random peptide sequences with length distributions identical to the MHC dataset. The random dataset yielded 483,448 and 600,624 unique 12-residue degenerate sequences for MHC I and II, respectively. In addition, we constructed a dictionary with random sequences with the same length-distribution as UniRef50.

### Word2Function

#### ExpGO65 dataset

Protein sequences with GO annotation were downloaded from the Gene Ontology website (https://geneontology.org/docs/download-go-annotations/)^49^. We filtered for sequences that have UniProt IDs and fall under the molecular function namespace. We then removed conflicting annotations for the same protein, resulting in a dataset of protein-GO term associations. 65 low-level GO terms were considered. The final dataset contains 56,299 protein sequences.

#### Supervised sequence-function model training

We use Protein Wordwise to parse an analyte protein into protein words. We generate embeddings for each residue by using ESM2. For each protein word, we compute the embedding by averaging the embeddings of the residues within the word. We treat the entire protein sequence as a single word and append its embedding to the end of the other word embeddings. Our supervised protein function prediction model is composed of Transformer^67^ layers and a fully connected layer. The model input consists of a CLS token embedding and protein word embeddings. The Transformer layers leverage self-attention mechanisms to capture interactions between words and aggregate all information into the CLS token. A fully connected layer then maps the representation of the CLS token to probabilities of function categories. The input embedding dimension and hidden layer dimension are both set to 1,280, with 20 attention heads. During training, we employed binary cross-entropy loss to optimize the model, using a batch size of 256,600 epochs, a weight decay of 1e-5, and an initial learning rate of 1e-3. Adam optimizer was used. To prevent overfitting, an early stopping strategy was applied after 60 epochs in conjunction with a cosine learning rate decay schedule.

#### Word-function mapping

We employ a supervised learning model to predict protein functions with protein word embeddings. If the model predicts one of the 65 functions for a query protein, we use the Integrated Gradients algorithm^51^ to quantify the contribution of each word to the prediction. Integrated Gradients is an interpretability technique that calculates the integral of the model’s output with respect to its inputs to quantify feature contributions. By integrating along a path from a baseline input (*e.g.*, an all-zero vector) to the input, it estimates the contribution of each feature. In our study, we treat each word in the protein sequence as a feature and compute its contribution to the model’s output. Word2Function ranks the word importance to the predicted GO term for each protein word based on its attribution score, and determines the functional annotation words by using a cumulative attribution score-based cutoff value α. Only protein words with a positive attribution score are ranked. A positive attribution score indicates that the input protein word positively contributed to the final prediction of GO terms. A cumulative attribution score is the sum of the positive attribution scores for all protein words in a sequence. We use a cutoff value α, defined as the cumulative attribution score multiplied by a factor of 0.5. If the sum of attribution scores for the top-N protein words has attained this alpha value, we retain the top-N protein words as functional annotation words, discarding the other protein words. The functional annotation words will be saved to our functional annotation word table called “WordTableGO65”.

#### WordTableGO65 for protein function prediction

WordTableGO65 was used for functional annotation at the whole-protein level (*i.e.*, not at the protein word-level). We use Protein Wordwise to parse an analyte sequence into protein words and use WordTableGO65 to find exact matches with these protein words. The corresponding GO term is then assigned to the protein word and the protein sequence.

#### Function prediction for DUFs

The InterProDUF dataset was compiled from Pfam entries annotated as DUFs from the InterPro database^2^. Specifically, sequences in UniRef50 containing Pfam domains annotated as DUFs in InterPro were included, resulting in 1,494,146 protein sequences. To predict function for these DUFs, we employed Protein Wordwise to parse the InterProDUF sequences into protein words. Subsequently, we utilized WordTableGO65 to identify exact matches for these protein words. The corresponding GO term was then assigned to both the protein word and the corresponding protein sequence.

### Metrics used for comparing with PROSITE

We compare our toolkit with PROSITE on two tasks:

1. Task1 is GO-term prediction at the whole-protein level. A functional MCC is defined as the MCC for a binary classification problem to determine whether a protein sequence is associated with a specific GO term. A functional MCC for each GO term is calculated based on true positives (TP), false positives (FP), true negatives (TN), and false negatives (FN) from predictions (Equation 1). The mean functional MCC for all 65 GO terms provides an overall measure of model performance.

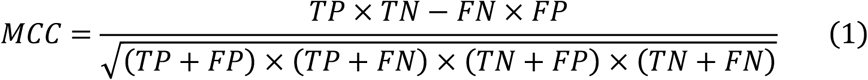
2. Task2 is functional annotation word/motif prediction. We use functional residue coverage as our metric for Protein Wordwise. We use word accuracy as our metric for Word2Function.

### PWNet

For all functional residue prediction tasks in the PWNet datasets, Pfam and GO information was obtained based on UniProt IDs, when available. The final PWNet datasets included proteins with lengths between 50 and 1,024 amino acids. An overview of these datasets are in Table S6. Data for protein-DNA and protein-RNA binding residues were sourced from the CLAPE dataset^68^, which was used for both training and testing. After merging and deduplication, the final dataset comprised 1,112 DNA-binding proteins and 991 RNA-binding proteins.

Data for small molecule- and metal ion-binding residues were obtained from Wang *et al.*, which contained 3,584 metal ion binding proteins and 4,964 small molecule binding proteins.

Data for ATP-binding residues was sourced from the E2EATP study^69^, utilizing the PATP-429 dataset, which consists of 429 sequences. Data for peptide-binding residues were derived from BioLiP2^70^, containing 39,065 initial sequences. After sequence retrieval based on UniProt IDs and data cleaning, 4,162 unique sequences remained. The enzyme active site dataset was obtained from the EasIFA study^42^, using the MCSA E-RXN CSA dataset with a total of 966 sequences.

Data for protein-protein interaction residues was sourced from the DIPS-Plus database. After converting structural annotations to sequence-based annotations and filtering, 5,851 homo-dimeric and 1,773 hetero-dimeric protein sequences were obtained. Data for Ion channel residues was compiled by ourselves. The names of ion channels and their subunits, along with their corresponding UniProt IDs, were retrieved from the IUPHAR/BPS Guide to PHARMACOLOGY (https://www.guidetopharmacology.org/)^71^. We obtained PDB structures, sequence lengths, and resolutions for each UniProt ID from UniProt, selecting a representative structure. Channels were identified using the Hole program, and residues within 10 Å of the channel center were labeled as channel-forming. A binary label was assigned to each residue, with “1” indicating a channel residue. All results were manually inspected and corrected, resulting in a final dataset of 95 valid entries.

## Data Availability

Our code is available at https://github.com/TianBoxue-lab/ProteinWordwise

## Acknowledgments

This work was supported by Beijing Frontier Research Center for Biological Structure (No. 041500002), CCF-Baidu Open Fund (2024), Tsinghua University Initiative Scientific Research Program (No.20231080030), and the Tsinghua-Peking University Center for Life Sciences (No.20111770319). We thank PaddlePaddle team for insightful technical discussions.

## Author Information

Correspondence and requests for materials should be addressed to B.T.

## Contributions

H.C. performed data analysis, figure preparation and attention head categorization. J.Z. developed Protein Wordwise. X.Z. assisted in the collection of PWNet datasets. J.C. developed Word2Function and generated WordTableGO65. L.G. and. X.X. helped to develop Protein Wordwise. X.L. assisted in analysis of GPCR words. X.L., X.Z. and B.T. revised the manuscript. B.T. designed the project, analyzed data, wrote and revised the manuscript.

## References

1 Chu, A. E., Lu, T. & Huang, P.-S. Sparks of function by *de novo* protein design. Nat. Biotechnol. 42, 203–215 (2024). 10.1038/s41587-024-02133-2

2 Paysan-Lafosse, T. et al. InterPro in 2022. Nucleic Acids Res. 51, D418–D427 (2023). 10.1093/nar/gkac993

3 Mistry, J. et al. Pfam: The protein families database in 2021. Nucleic Acids Res. 49, D412–D419 (2020). 10.1093/nar/gkaa913

4 Wang, X. et al. Multi-modal deep learning enables efficient and accurate annotation of enzymatic active sites. Nat. Commun. 15, 7348 (2024). 10.1038/s41467-024-51511-6

5 Yaffe, M. B. et al. A motif-based profile scanning approach for genome-wide prediction of signaling pathways. Nat. Biotechnol. 19, 348–353 (2001). 10.1038/86737

6 Fowler, D. M. & Fields, S. Deep mutational scanning: a new style of protein science. Nat. Methods 11, 801–807 (2014). 10.1038/nmeth.3027

7 Nagarajan, V. & Elasri, M. O. Structure and function predictions of the Msa protein in *Staphylococcus aureus*. BMC bioinform. 8, 1–9 (2007). 10.1186/1471-2105-8-S7-S5

8 Sigrist, C. J. et al. PROSITE, a protein domain database for functional characterization and annotation. Nucleic Acids Res. 38, D161–D166 (2010). 10.1093/nar/gkp885

9 Consortium, T. U. UniProt: the universal protein knowledgebase. Nucleic Acids Res. 45, D158–D169 (2017). 10.1093/nar/gkw1099

10 Boeckmann, B. et al. The SWISS-PROT protein knowledgebase and its supplement TrEMBL in 2003. Nucleic Acids Res. 31, 365–370 (2003). 10.1093/nar/gkg095

11 Notin, P. et al. Proteingym: Large-scale benchmarks for protein fitness prediction and design. Adv. Neural Inf. Process. Syst.36, 64331–64379 (2024).

12 Chen, X. et al. Integrative residue-intuitive machine learning and MD Approach to Unveil Allosteric Site and Mechanism for β_2_AR. Nat. Commun. 15, 8130 (2024). 10.1038/s41467-024-52399-y

13 Cheng, Y. et al. Co-evolution-based prediction of metal-binding sites in proteomes by machine learning. Nat. Chem. Biol. 19, 548–555 (2023). 10.1038/s41589-022-01223-z

14 Cagiada, M. et al. Discovering functionally important sites in proteins. Nat. Commun. 14, 4175 (2023). 10.1038/s41467-023-39909-0

15 Jang, Y. J. et al. Accurate prediction of protein function using statistics-informed graph networks. Nat. Commun. 15, 6601 (2024). 10.1038/s41467-024-50955-0

16 Chen, Y., Xu, Y., Liu, D., Xing, Y. & Gong, H. An end-to-end framework for the prediction of protein structure and fitness from single sequence. Nat. Commun. 15, 7400 (2024). 10.1038/s41467-024-51776-x

17 Ma, Z. et al. EvoAI enables extreme compression and reconstruction of the protein sequence space. Nat. Methods 22, 102–112 (2024). 10.1038/s41592-024-02504-2

18 Lin, Z. et al. Evolutionary-scale prediction of atomic-level protein structure with a language model. Science 379, 1123–1130 (2023). 10.1126/science.ade2574

19 Hayes, T. et al. Simulating 500 million years of evolution with a language model. Science 0, eads0018 (2025). 10.1126/science.ads0018

20 Brandes, N., Ofer, D., Peleg, Y., Rappoport, N. & Linial, M. ProteinBERT: a universal deep-learning model of protein sequence and function. Bioinformatics 38, 2102–2110 (2022). 10.1093/bioinformatics/btac020

21 Vu, M. H. et al. Linguistically inspired roadmap for building biologically reliable protein language models. *Nat*. Mach. Intell. 5, 485–496 (2023). 10.1038/s42256-023-00637-1

22 Unsal, S. et al. Learning functional properties of proteins with language models. *Nat*. Mach. Intell. 4, 227–245 (2022). 10.1038/s42256-022-00457-9

23 Biswas, S., Khimulya, G., Alley, E. C., Esvelt, K. M. & Church, G. M. Low-N protein engineering with data-efficient deep learning. Nat. Methods 18, 389–396 (2021). 10.1038/s41592-021-01100-y

24 Guo, L., et al. Enhancing Enzyme Activity with Mutation Combinations Guided by Few-shot Learning and Causal Inference. Preprint at Res. Sq. 10.21203/rs.3.rs-5354708/v1 (2024).

25 Yu, T. et al. Enzyme function prediction using contrastive learning. Science 379, 1358–1363 (2023). 10.1126/science.adf2465

26 Alva, V., Söding, J. & Lupas, A. N. A vocabulary of ancient peptides at the origin of folded proteins. eLife 4, e09410 (2015). 10.7554/eLife.09410

27 de Juan, D., Pazos, F. & Valencia, A. Emerging methods in protein co-evolution. Nat. Rev. Genet. 14, 249–261 (2013). 10.1038/nrg3414

28 Wang, J., Liu, Y. & Tian, B. Protein-small molecule binding site prediction based on a pre-trained protein language model with contrastive learning. J. Cheminf. 16, 125 (2024). 10.1186/s13321-024-00920-2

29 Kulmanov, M. et al. Protein function prediction as approximate semantic entailment. *Nat*. Mach. Intell. 6, 220–228 (2024). 10.1038/s42256-024-00795-w

30 Kudo, T. Subword regularization: Improving neural network translation models with multiple subword candidates. in Proceedings of the 56th Annual Meeting of the Association for Computational Linguistics. (Volume 1: Long Papers) 2018; 66–75

31 Osmanbeyoglu, H. U. & Ganapathiraju, M. K. N-gram analysis of 970 microbial organisms reveals presence of biological language models. BMC Bioinform. 12, 1–12 (2011). 10.1186/1471-2105-12-12

32 Mandal, S. & Bhattacharyya, D. Two L-amino acid oxidase isoenzymes from Russell’s viper (*Daboia russelli russelli*) venom with different mechanisms of inhibition by substrate analogs. FEBS J. 275, 2078–2095 (2008). 10.1111/j.1742-4658.2008.06362.x

33 Jha, K., Saha, S. & Singh, H. Prediction of protein-protein interaction using graph neural networks. Sci. Rep. 12, 8360 (2022). 10.1038/s41598-022-12201-9

34 Blondel, V. D., Guillaume, J.-L., Lambiotte, R. & Lefebvre, E. Fast unfolding of communities in large networks. J. Stat. Mech. Theory Exp. 2008, P10008 (2008). 10.1088/1742-5468/2008/10/P10008

35 Heydenreich, F. M. et al. Molecular determinants of ligand efficacy and potency in GPCR signaling. Science 382, eadh1859 (2023). 10.1126/science.adh1859

36 Suzek, B. E. et al. UniRef clusters: a comprehensive and scalable alternative for improving sequence similarity searches. Bioinformatics 31, 926–932 (2015). 10.1093/bioinformatics/btu739

37 Schulze, T. K. & Lindorff-Larsen, K. Effects of residue substitutions on the cellular abundance of proteins. eLife 13, RP103721 (2024). 10.7554/elife.103721.1

38 Gilston, B. A. et al. Structural and mechanistic basis of zinc regulation across the *E. coli* Zur regulon. PLoS Biol. 12, e1001987 (2014). 10.1371/journal.pbio.1001987

39 Fuhrmann, J. et al. McsB is a protein arginine kinase that phosphorylates and inhibits the heat-shock regulator CtsR. Science 324, 1323–1327 (2009). 10.1126/science.1170088

40 Chan, K. H. & Wong, K. B. Structure of an essential GTPase, YsxC, from Thermotoga maritima. Acta Crystallogr. Sect. F-Struct. Biol. Cryst. Commun. 67, 640–646 (2011). 10.1107/S1744309111011651

41 Tahirov, T. H. et al. High-resolution Crystal Structures of Two Polymorphs of Cytochrome *c*’ from the Purple Phototrophic Bacterium *Rhodobacter capsulatus*. J. Mol. Biol. 259, 467–479 (1996). 10.1006/jmbi.1996.0333

42 Wang, X. et al. Multi-modal deep learning enables efficient and accurate annotation of enzymatic active sites. Nat. Commun. 15, 7348 (2024). 10.1038/s41467-024-51511-6

43 Dai, S., Schwendtmayer, C., Schurmann, P., Ramaswamy, S. & Eklund, H. Redox signaling in chloroplasts: cleavage of disulfides by an iron-sulfur cluster. Science 287, 655–658 (2000). 10.1126/science.287.5453.655

44 Hart, P. J. et al. A structure-based mechanism for copper-zinc superoxide dismutase. Biochemistry 38, 2167–2178 (1999). 10.1021/bi982284u

45 Morehead, A., Chen, C., Sedova, A. & Cheng, J. DIPS-Plus: The enhanced database of interacting protein structures for interface prediction. Sci. Data 10, 509 (2023). 10.1038/s41597-023-02409-3

46 Wu, L. C., Tuot, D. S., Lyons, D. S., Garcia, K. C. & Davis, M. M. Two-step binding mechanism for T-cell receptor recognition of peptide-MHC. Nature 418, 552–556 (2002). 10.1038/nature00920

47 Tadros, D. M., Eggenschwiler, S., Racle, J. & Gfeller, D. The MHC Motif Atlas: a database of MHC binding specificities and ligands. Nucleic Acids Res. 51, D428–D437 (2023). 10.1093/nar/gkac965

48 Dönnes, P. & Elofsson, A. Prediction of MHC class I binding peptides, using SVMHC. BMC Bioinform. 3, 25 (2002). 10.1186/1471-2105-3-25

49 Ashburner, M. et al. Gene Ontology: tool for the unification of biology. Nat. Genet. 25, 25–29 (2000). 10.1038/75556

50 Jang, Y. J. et al. Accurate prediction of protein function using statistics-informed graph networks. Nat. Commun. 15, 6601 (2024). 10.1038/s41467-024-50955-0

51 Sundararajan, M., Taly, A. & Yan, Q. Axiomatic attribution for deep networks. in Proceedings of the 34th International Conference on Machine Learning. 70, 3319–3328 (JMLR.org, Sydney, NSW, Australia, 2017).

52 Minajigi, A. & Francklyn, C. S. RNA-assisted catalysis in a protein enzyme: The 2’-hydroxyl of tRNA (Thr) A76 promotes aminoacylation by threonyl-tRNA synthetase. Proc. Natl Acad. Sci. USA 105, 17748–17753 (2008). 10.1073/pnas.0804247105

53 Guerra, D. J. & Browse, J. A. *Escherichia coli* β-hydroxydecanoyl thioester dehydrase reacts with native C_10_ acyl-acyl-carrier proteins of plant and bacterial origin. Arch. Biochem. Biophys. 280, 336–345 (1990). 10.1016/0003-9861(90)90339-Z

54 Jeltsch, A. The Cytosine N^4^-Methyltransferase M.*Pvu*II Also Modifies Adenine Residues. Biol. Chem. 382, 707–710 (2001). 10.1515/BC.2001.084

55 Nakatani, T. et al. Enhancement of thioredoxin/glutaredoxin-mediated L-cysteine synthesis from S-sulfocysteine increases L-cysteine production in *Escherichia coli*. Microb. Cell Fact. 11, 62 (2012). 10.1186/1475-2859-11-62

56 Fonfría-Subirós, E. et al. Crystal structure of a complex of DNA with one AT-hook of HMGA1. PLoS One 7, e37120 (2012). 10.1371/journal.pone.0037120

57 Pereira, B. et al. CDX2 regulation by the RNA-binding protein MEX3A: impact on intestinal differentiation and stemness. Nucleic Acids Res. 41, 3986–3999 (2013). 10.1093/nar/gkt087

58 Inoue, M. et al. Nucleotide-dependent conformational changes and assembly of the AAA ATPase SKD1/VPS4B. Traffic 9, 2180–2189 (2008). 10.1111/j.1600-0854.2008.00831.x

59 Zhang, X. et al. Assignment of function to a domain of unknown function: DUF1537 is a new kinase family in catabolic pathways for acid sugars. Proc. Natl Acad. Sci. USA 113, E4161–4169 (2016). 10.1073/pnas.1605546113

60 Yuan, Y. et al. Chemoproteomic discovery of a human RNA ligase. Nat. Commun. 14, 842 (2023). 10.1038/s41467-023-36451-x

61 Li, H. J. et al. POD1 regulates pollen tube guidance in response to micropylar female signaling and acts in early embryo patterning in *Arabidopsis*. Plant Cell 23, 3288–3302 (2011). 10.1105/tpc.111.088914

62 Nada, H. et al. New insights into protein-protein interaction modulators in drug discovery and therapeutic advance. Sig. Transduct. Target Ther. 9, 341 (2024). 10.1038/s41392-024-02036-3

63 Souza, P. C. T. et al. Martini 3: a general purpose force field for coarse-grained molecular dynamics. Nat. Methods 18, 382–388 (2021). 10.1038/s41592-021-01098-3

64 Sanabria, M., Hirsch, J., Joubert, P. M. & Poetsch, A. R. DNA language model GROVER learns sequence context in the human genome. *Nat*. Mach. Intell. 6, 911–923 (2024). 10.1038/s42256-024-00872-0

65 Shen, T. et al. Accurate RNA 3D structure prediction using a language model-based deep learning approach. Nat. Methods 21, 2287–2298 (2024). 10.1038/s41592-024-02487-0

66 Hao, M. et al. Large-scale foundation model on single-cell transcriptomics. Nat. Methods 21, 1481–1491 (2024). 10.1038/s41592-024-02305-7

## References

67 Vaswani, A. et al. Attention is All you Need. in Advances in Neural Information Processing Systems (eds. Guyon, I. et al.) 30 (Curran Associates, Inc., 2017).

68 Liu, Y. & Tian, B. Protein-DNA binding sites prediction based on pre-trained protein language model and contrastive learning. Brief. Bioinform. 25 (2023). 10.1093/bib/bbad488

69 Rao, B., Yu, X., Bai, J. & Hu, J. E2EATP: Fast and High-Accuracy Protein-ATP Binding Residue Prediction via Protein Language Model Embedding. J. Chem. Inf. Model. 64, 289–300 (2024). 10.1021/acs.jcim.3c01298

70 Zhang, C., Zhang, X., Freddolino, P. L. & Zhang, Y. BioLiP2: an updated structure database for biologically relevant ligand-protein interactions. Nucleic Acids Res. 52, D404–D412 (2024). 10.1093/nar/gkad630

71 Harding, S. D. et al. The IUPHAR/BPS Guide to PHARMACOLOGY in 2024. Nucleic Acids Res. 52, D1438–D1449 (2023). 10.1093/nar/gkad944

