## Supplementary figures and images for "Automatically Defining Protein Words for Diverse Functional Predictions Based on Attention Analysis of a Protein Language Model"

### Supplemental Figure1

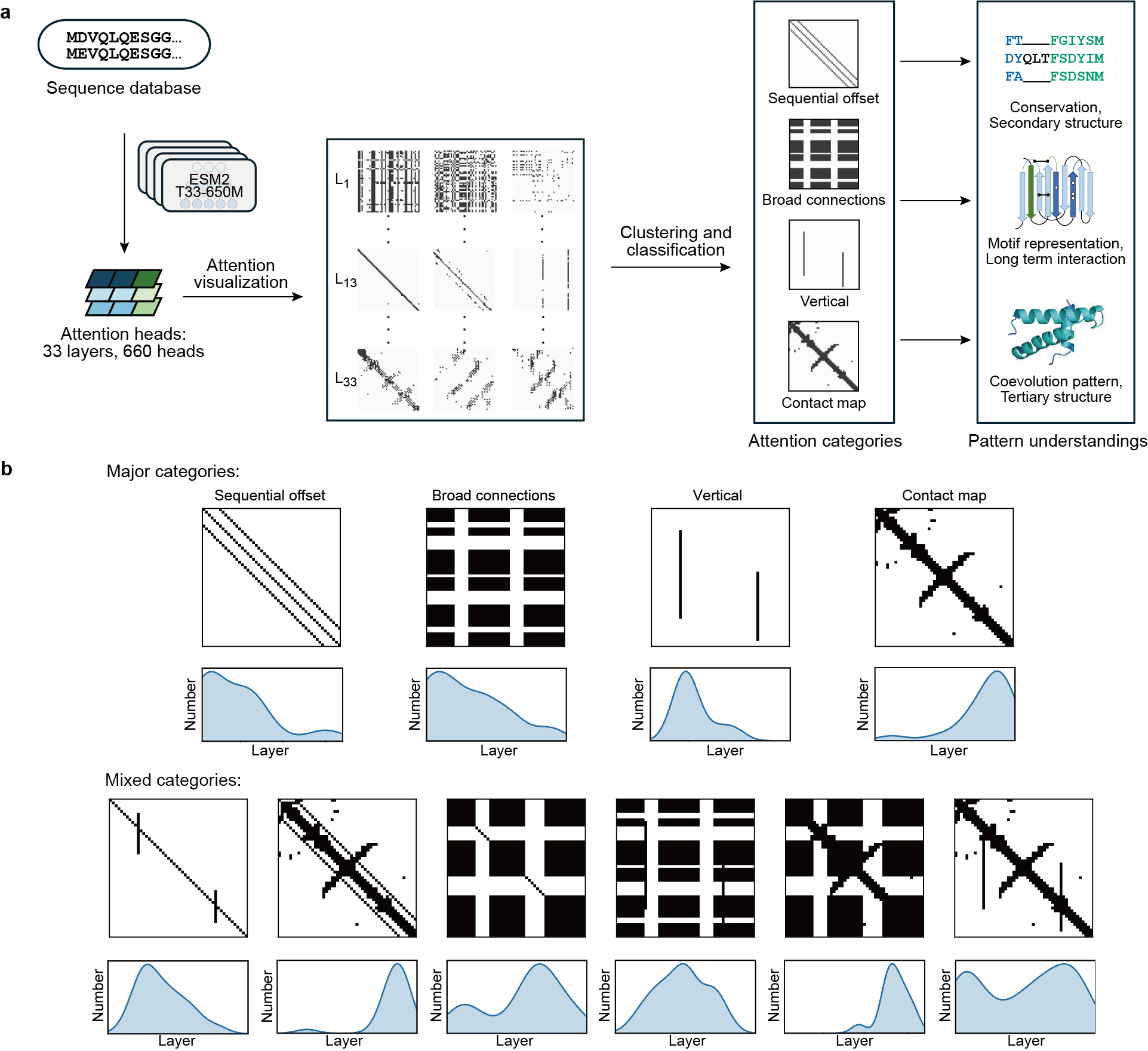

### Supplemental Figure2

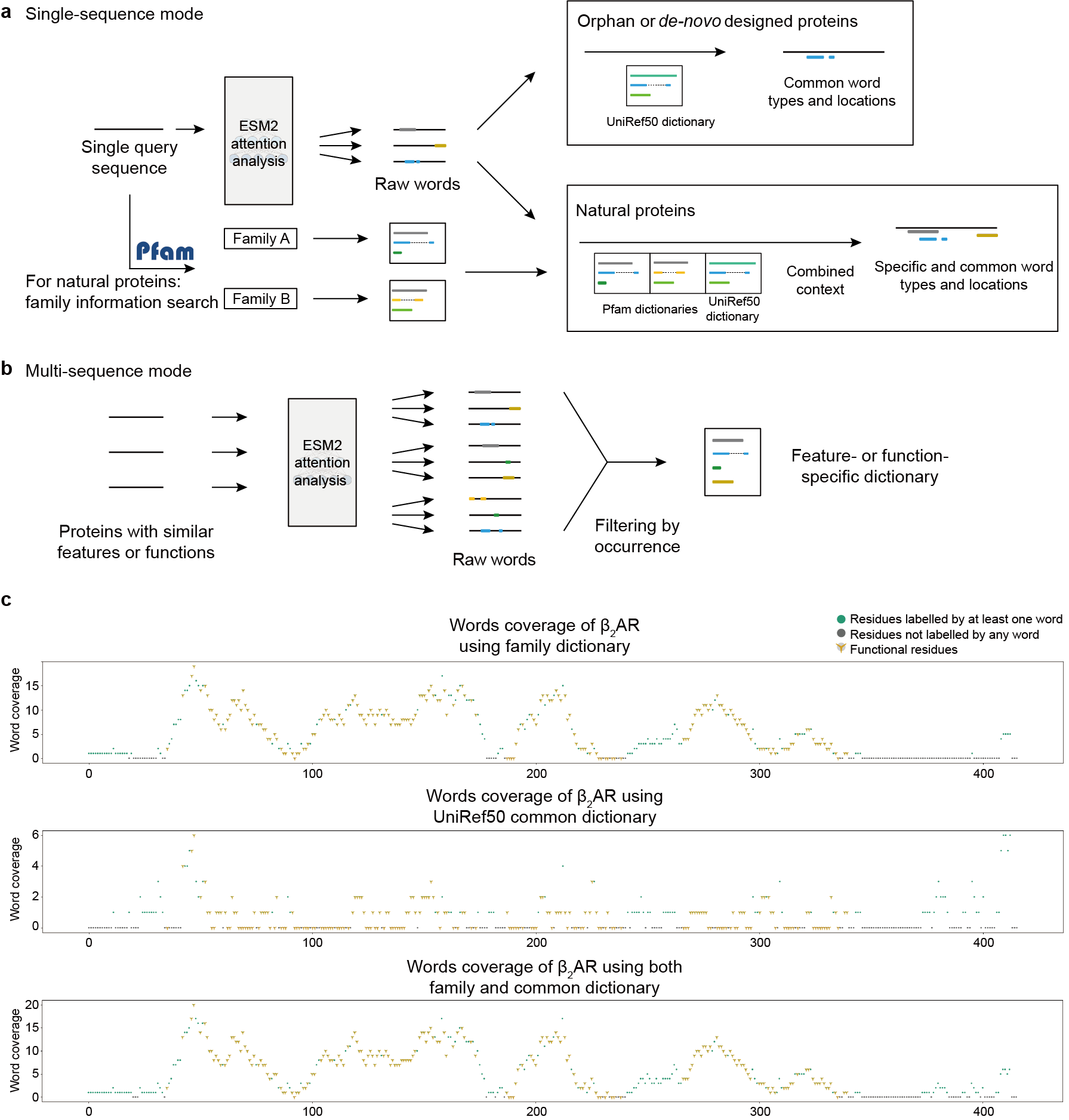

### Supplemental Figure3

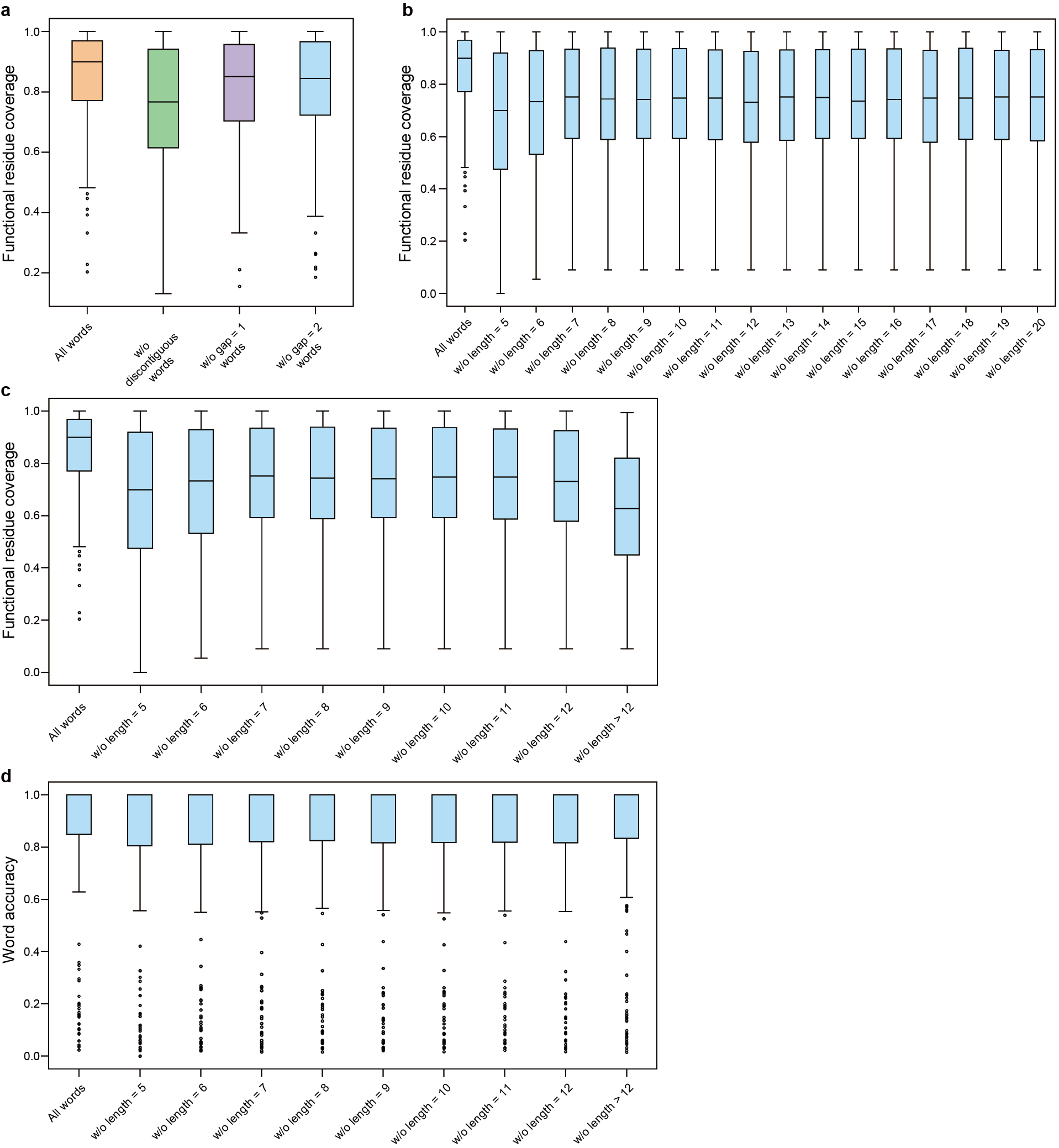

### Supplemental Figure4

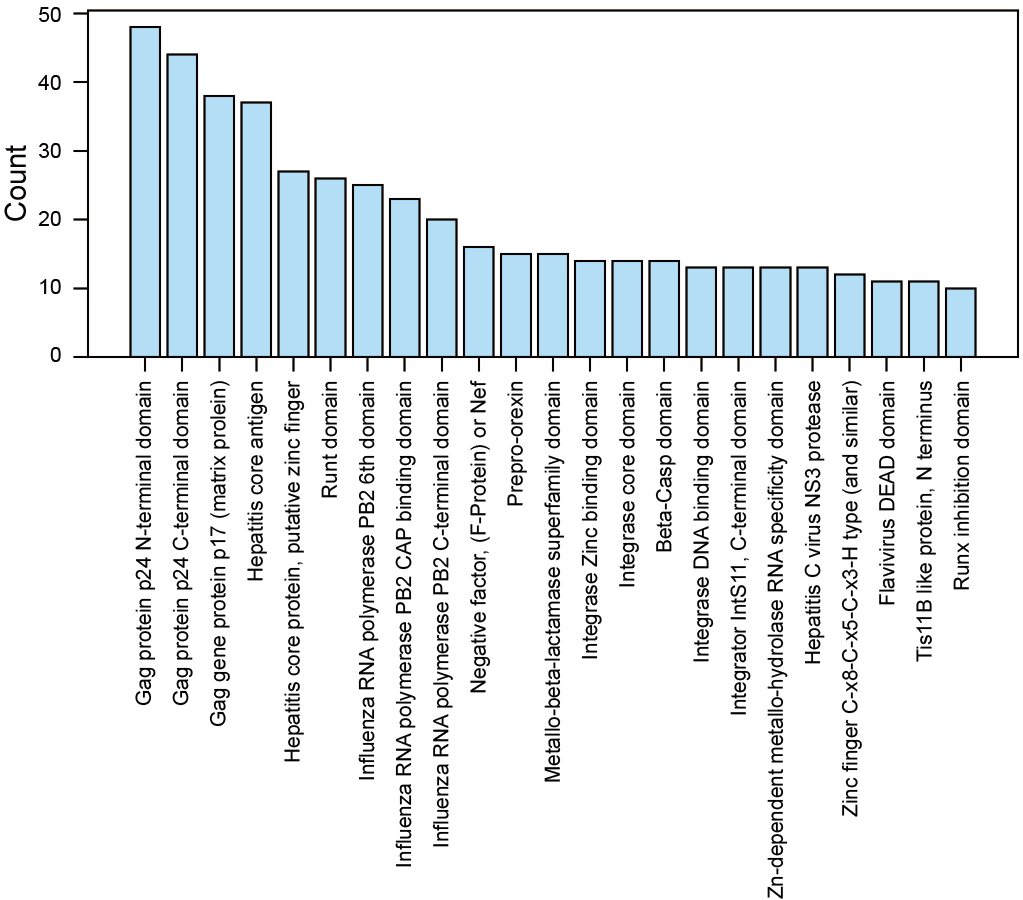
