## Supplemental Table1-9 for "Automatically Defining Protein Words for Diverse Functional Predictions Based on Attention Analysis of a Protein Language Model"

### Supplementary Information Guide

#### List of Supplementary Figures

Supplementary Figure 1: Attention categories of the PLM (ESM2).
Supplementary Figure 2: Single- and multi-sequence modes of Protein Wordwise.
Supplementary Figure 3: Ablation studies of Protein Wordwise using the DMS dataset.
Supplementary Figure 4: Distribution of Pfam domains among the 1,202 proteins containing MHC peptides and having experimental 3D structures.

#### List of Supplementary Tables

Supplementary Table 1: Tunning hyperparameters based on the $\beta_2$AR experimental data.
Supplementary Table 2: The 12-degenerated residue types defined in this study.
Supplementary Table 3: Functional residue coverage and word accuracy comparison among different schemes for residue degeneration for $\beta_2$AR.
Supplementary Table 4: Tunning of schemes for residue degeneration based on the $\beta_2$AR experimental data.
Supplementary Table 5: A list of 16 DMS dataset proteins for which Protein Wordwise achieved 100% predicted functional residue coverage.
Supplementary Table 6: An overview of datasets comprising the PWNet data resource for functional residue prediction.
Supplementary Table 7: GO terms used in the ExpGO65 dataset.
Supplementary Table 8: Prediction performance (functional MCCs) for Word2Function for ExpGO65.
Supplementary Table 9: Prediction performance (functional MCCs) for WordTableGO65 for ExpGO65.

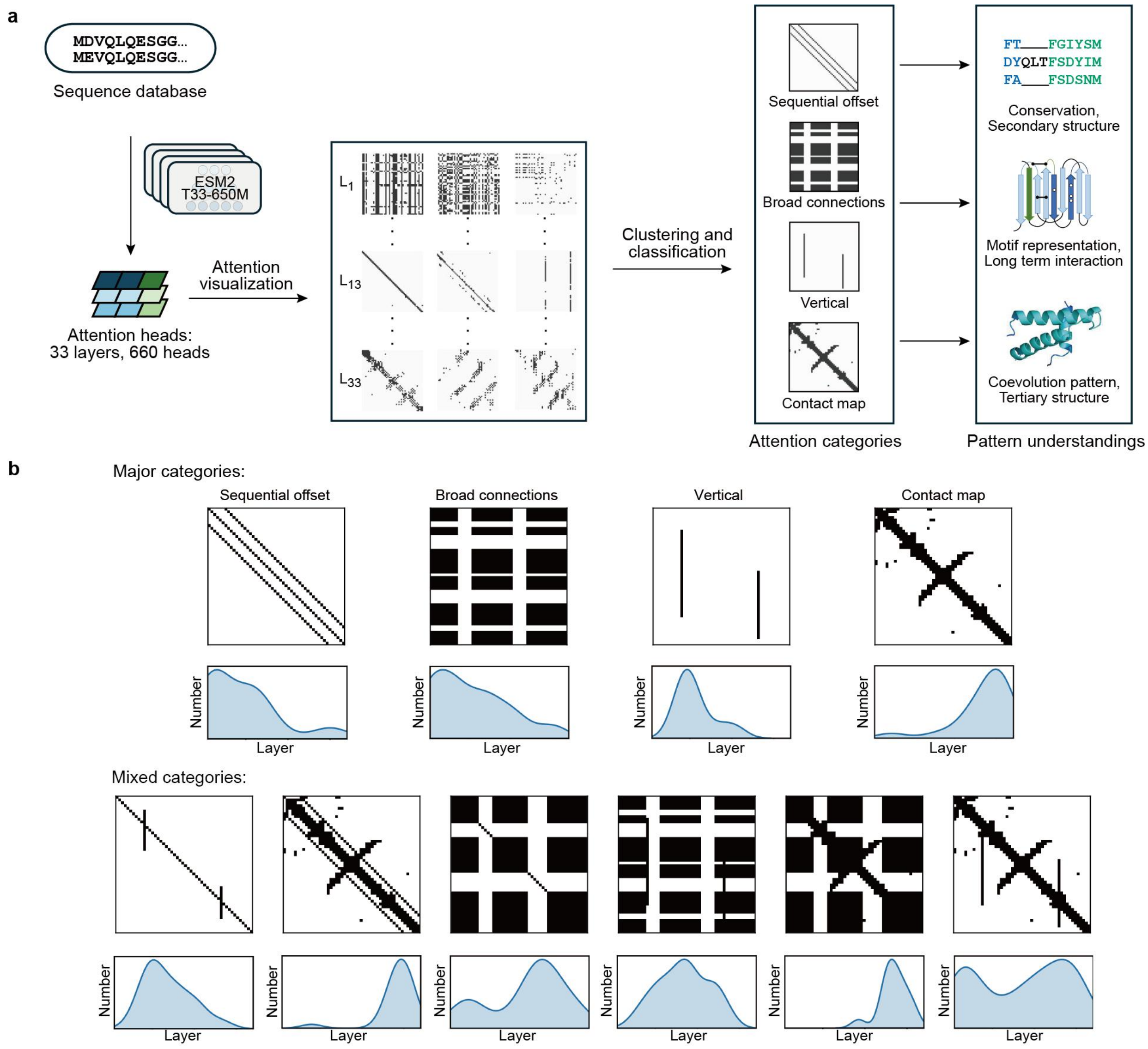


**Supplementary Figure 1 | Attention categories of the PLM (ESM2). a**, Four major attention head categories. In the first step, the protein sequence is passed to ESM2 T33-650M model for attention prediction. Based on the observations of from >1,000 sequences, we identified 4 major attention categories: sequential offset, broad connections, vertical (key residues) and contact map. The categories of attention heads were consistent across different input sequences. **b**, An overview of all attention head categories. The top images depict the categories and the bottom images represent the distribution of the attention heads for the same category across different layers.

The contact map categories are mostly located in Layers 31-33; the broad connection categories are mostly located in Layers 1-10.

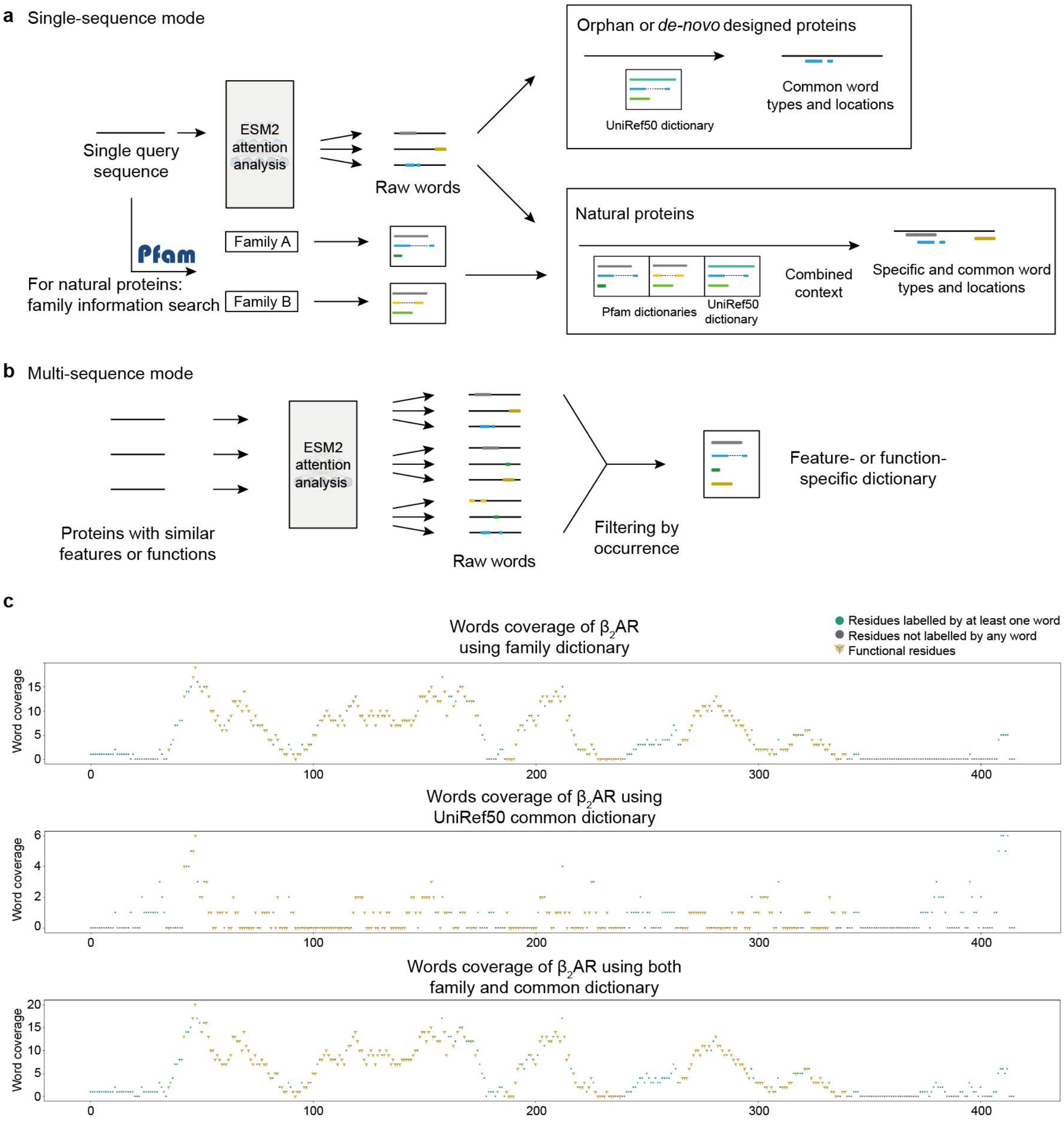


**Supplementary Figure 2 | Single- and multi-sequence modes of Protein Wordwise. a,** Flowchart for the single-sequence mode of Protein Wordwise. For an analyte protein sequence lacking Pfam information, protein words were predicted by matching raw words predicted by the Louvain algorithm and the UniRef50 dictionary. For analyte sequences having known UniProt ID and Pfam, the UniRef50 and all suitable Pfam dictionaries are used for matching. **b**, Flowchart for the multi-sequence mode of Protein Wordwise. Protein words are predicted from all sequences using the Louvain algorithm, stratified by length; only protein words with the highest occurrences (for each length bin) are selected for dictionary construct ion (Methods). **c**, An example of functional residue coverage and functional residue prediction using solely a Pfam dictionary (Pfam PF00001), using solely a UniRef50 dictionary, or using both dictionaries.

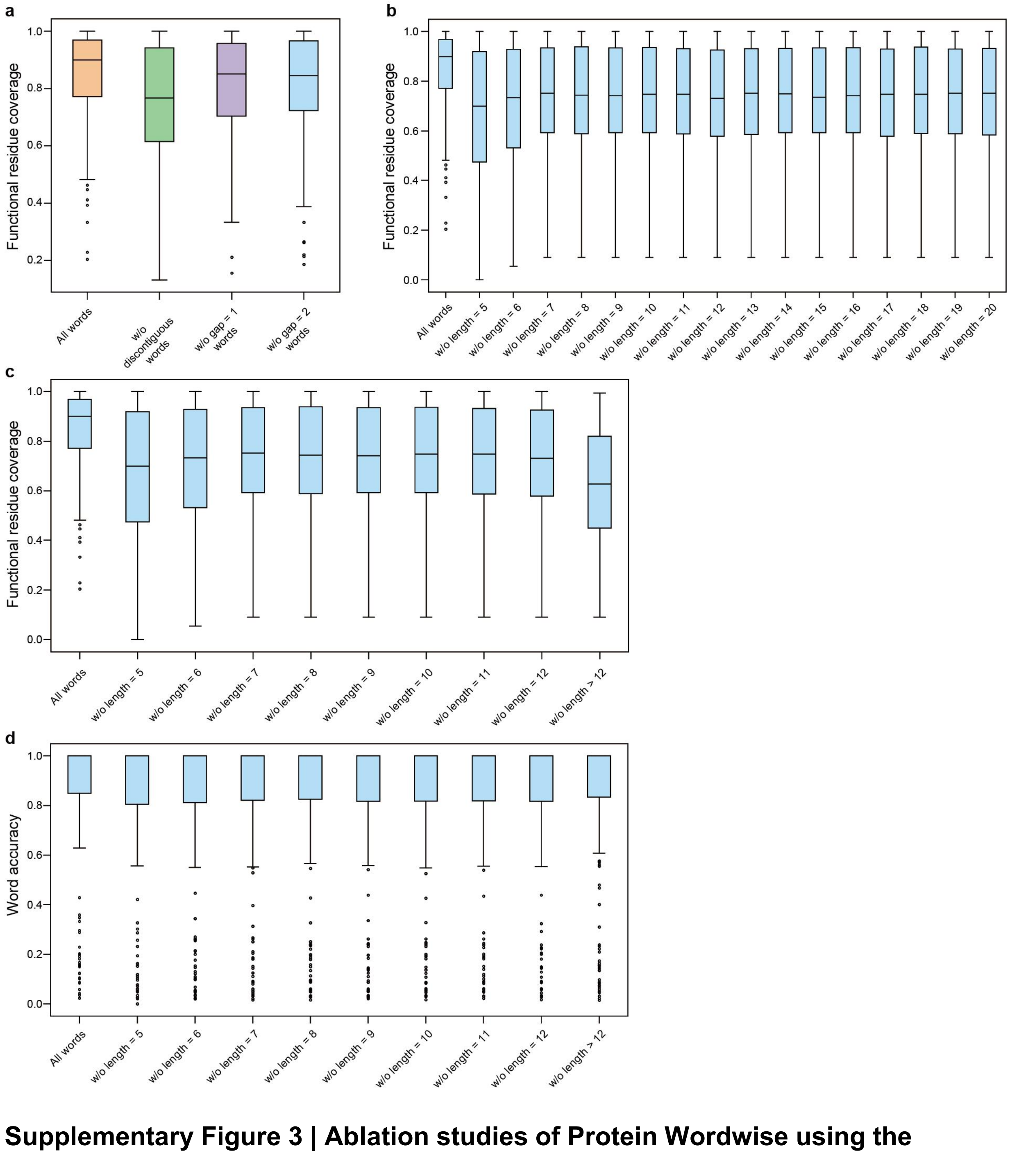


**Supplementary Figure 3 | Ablation studies of Protein Wordwise using the DMS dataset.** **a,** Functional residue coverage performance of Protein Wordwise after removing discontiguous words, discontiguous words with one gap, or discontiguous words with two gaps. **b,** Functional residue coverage performance of Protein Wordwise after removing words of differing lengths as indicated. **c,** After merging words of length greater than 12, the functional residue coverage performance of Protein Wordwise upon removing words with different lengths. **d,** The accuracy of Protein Wordwise upon removing words with different lengths in the DMS dataset.

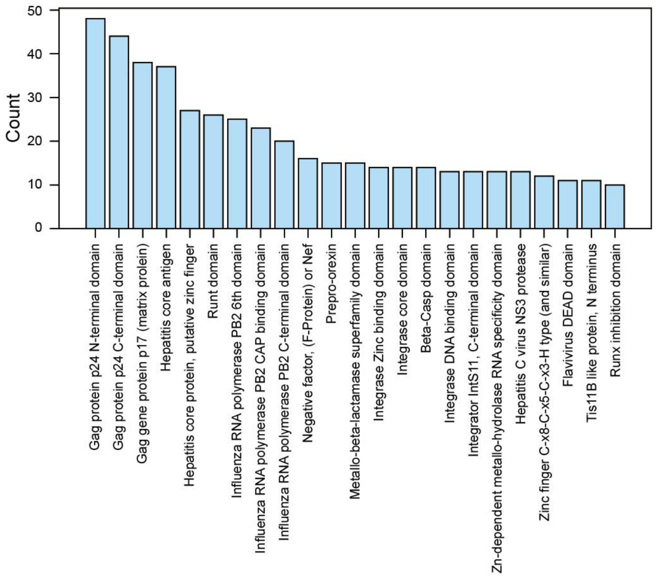


**Supplementary Figure 4 | Distribution of Pfam domains among the 1,202 proteins containing MHC peptides and having experimental 3D structures.** The top three Pfam domains were all Gag, with 48, 42, and 38 proteins, respectively. Gag is a structural protein found in HIV-1 and other retroviruses.

**Supplementary Table 1 | Tunning hyperparameters based on the $\beta_2$AR experimental data.**

| Contiguous short word percentage* | Contiguous long word percentage* | Discontiguous short word percentage* | Discontiguous long word percentage* | Number of words | Functional residue coverage | Word accuracy |
|---|---|---|---|---|---|---|
| 0.025 | 0.0025 | 0.000 | 0.0000 | 108 | 0.908 | 0.907 |
| 0.025 | 0.0050 | 0.000 | 0.0000 | 128 | 0.923 | 0.852 |
| 0.025 | 0.0075 | 0.000 | 0.0000 | 161 | 0.954 | 0.745 |
| 0.050 | 0.0025 | 0.000 | 0.0000 | 113 | 0.908 | 0.903 |
| 0.050 | 0.0050 | 0.000 | 0.0000 | 133 | 0.923 | 0.850 |
| 0.050 | 0.0075 | 0.000 | 0.0000 | 166 | 0.954 | 0.747 |
| 0.050 | 0.0100 | 0.000 | 0.0000 | 190 | 0.969 | 0.726 |
| 0.050 | 0.0150 | 0.000 | 0.0000 | 230 | 0.990 | 0.683 |
| 0.050 | 0.0200 | 0.000 | 0.0000 | 250 | 1.000 | 0.660 |
| 0.050 | 0.0250 | 0.000 | 0.0000 | 262 | 1.000 | 0.653 |
| 0.050 | 0.0300 | 0.000 | 0.0000 | 269 | 1.000 | 0.651 |
| 0.075 | 0.0025 | 0.000 | 0.0000 | 119 | 0.908 | 0.874 |
| 0.075 | 0.0050 | 0.000 | 0.0000 | 139 | 0.923 | 0.827 |
| 0.075 | 0.0075 | 0.000 | 0.0000 | 172 | 0.954 | 0.733 |
| 0.150 | 0.0100 | 0.000 | 0.0000 | 204 | 0.969 | 0.711 |
| 0.150 | 0.0150 | 0.000 | 0.0000 | 244 | 0.990 | 0.672 |
| 0.150 | 0.0200 | 0.000 | 0.0000 | 264 | 1.000 | 0.652 |
| 0.150 | 0.0250 | 0.000 | 0.0000 | 276 | 1.000 | 0.645 |
| 0.150 | 0.0300 | 0.000 | 0.0000 | 283 | 1.000 | 0.643 |
| 0.100 | 0.0100 | 0.000 | 0.0000 | 200 | 0.969 | 0.715 |
| 0.100 | 0.0150 | 0.000 | 0.0000 | 240 | 0.990 | 0.675 |
| 0.100 | 0.0200 | 0.000 | 0.0000 | 260 | 1.000 | 0.654 |
| 0.100 | 0.0250 | 0.000 | 0.0000 | 272 | 1.000 | 0.647 |
| 0.100 | 0.0300 | 0.000 | 0.0000 | 279 | 1.000 | 0.645 |
| 0.025 | 0.0025 | 0.015 | 0.0010 | 167 | 0.959 | 0.928 |
| 0.025 | 0.0025 | 0.015 | 0.0020 | 186 | 0.979 | 0.930 |
| 0.025 | 0.0025 | 0.010 | 0.0025 | 191 | 0.985 | 0.927 |
| 0.025 | 0.0025 | 0.025 | 0.0010 | 174 | 0.969 | 0.931 |

| | | | | | | |
|---|---|---|---|---|---|---|
| **0.025** | **0.0025** | **0.025** | **0.0015** | **183** | **0.985** | **0.929** |
| 0.025 | 0.0025 | 0.025 | 0.0020 | 193 | 0.985 | 0.933 |
| 0.025 | 0.0025 | 0.025 | 0.0025 | 201 | 0.990 | 0.930 |
| 0.025 | 0.0025 | 0.025 | 0.0050 | 232 | 0.990 | 0.922 |
| 0.025 | 0.0025 | 0.025 | 0.0075 | 259 | 0.990 | 0.896 |
| 0.025 | 0.0025 | 0.025 | 0.0100 | 273 | 0.990 | 0.894 |
| 0.025 | 0.0025 | 0.020 | 0.0015 | 177 | 0.979 | 0.927 |
| 0.025 | 0.0025 | 0.020 | 0.0020 | 187 | 0.979 | 0.930 |
| 0.025 | 0.0025 | 0.020 | 0.0025 | 195 | 0.985 | 0.928 |
| 0.025 | 0.0025 | 0.100 | 0.0025 | 228 | 0.995 | 0.899 |
| 0.025 | 0.0025 | 0.100 | 0.0050 | 259 | 0.995 | 0.896 |
| 0.025 | 0.0025 | 0.100 | 0.0075 | 286 | 0.995 | 0.874 |

* Length for short words ranges from 5 to 6; length for long words ranges from 7 to 20. The UniRef50 dictionary was not included in these predictions.

**Supplementary Table 2 | The 12-degenerated residue types defined in this study**

| Residue type | One code amino acid | Notes |
|---|---|---|
| A | D,E | Negatively charged |
| B | K,R | Positively charged |
| C | C | Cysteine |
| H | L,I,V | Hydrophobic (branched chain) |
| H | H | Histidine |
| M | M | Methionine |
| N | N,Q | Amino acids with an amide group |
| O | S,T | Amino acids with hydroxyl |
| S | A,G | Hydrophobic (short chain) |
| P | P | Proline |
| Y | Y | Tyrosine |
| @ | F,W | Aromatic |

**Supplementary Table 3 | Functional residue coverage and word accuracy comparison among different schemes for residue degeneration for $\beta_2$AR**

| Degenerate residue scheme | Functional residue coverage | Word accuracy |
|---|---|---|
| 4 | 0.918 | 0.752 |
| **12** | **0.985** | **0.929** |
| 20 | 0.918 | 0.862 |

The UniRef50 dictionary was not included in these predictions.

**Supplementary Table 4 | Tunning of schemes for residue degeneration based on the $\beta_2$AR experimental data.**

| Residue | 4-type codes | Remarks | 12-type codes | Remarks |
|---|---|---|---|---|
| D | a | Negatively charged | a | Negatively charged |
| E | | | | |
| K | b | Positively charged | b | Positively charged |
| R | | | | |
| H | | | H | Histidine |
| A | h | Hydrophobic | s | Hydrophobic, with small sidechain |
| G | | | | |
| I | | | h | Hydrophobic |
| L | | | | |
| V | | | | |
| M | | | M | Methionine |
| P | | | P | Proline |
| F | | | @ | Aromatic |
| W | | | | |
| S | p | Polar | o | Polar, with hydroxyl |
| T | | | | |
| N | | | n | Amides |
| Q | | | | |
| Y | | | Y | Tyrosine |
| C | | | C | Cysteine |

**Supplementary Table 5 | A list of 16 DMS dataset proteins for which Protein Wordwise achieved 100% predicted functional residue coverage**

| Protein name | UniProt ID | Sequence length | Number of functional residues / Sequence length | Number of predicted residues / Sequence length |
|---|---|---|---|---|
| A4_HUMAN_ (Seuma_2022) | P05067 | 770 | 0.055 | 0.983 |
| BLAT_ECOLX_ (Firnberg_2014) | P62593 | 286 | 0.832 | 1.000 |
| CCDB_ECOLI_ (Adkar_2012) | P62554 | 101 | 0.505 | 0.931 |
| DNJA1_HUMAN_ (Tsuboyama_2023) | P31689 | 65 | 1.000 | 1.000 |
| DYR_ECOLI_ (Thompson_2019) | P0ABQ4 | 159 | 0.925 | 1.000 |
| GAL4_YEAST_ (Kitzman_2015) | P04386 | 881 | 0.073 | 0.839 |
| TADBP_HUMAN_ (Bolognesi_2019) | Q13148 | 414 | 0.200 | 0.995 |
| MK01_HUMAN_ (Brenan_2016) | P28482 | 360 | 0.939 | 1.000 |
| PTEN_HUMAN_ (Matreyek_2021) | P60484 | 403 | 0.868 | 1.000 |
| PTEN_HUMAN_ (Mighell_2018) | P60484 | 403 | 0.988 | 1.000 |
| RL40A_YEAST_ (Roscoe_2013) | P0CH08 | 128 | 0.531 | 1.000 |
| RL40A_YEAST_ (Roscoe_2014) | P0CH08 | 128 | 0.555 | 1.000 |
| SRC_HUMAN_ (Ahler_2019) | P12931 | 536 | 0.407 | 0.953 |
| SRC_HUMAN_ (Chakraborty_2023) | P12931 | 536 | 0.405 | 0.924 |
| SRC_HUMAN_ (Nguyen_2022) | P12931 | 536 | 0.388 | 0.924 |
| UBC9_HUMAN_ (Weile_2017) | P63279 | 159 | 0.893 | 0.994 |

**Supplementary Table 6 | An overview of datasets comprising the PWNet data resource for functional residue prediction**

| Dataset | Average label ratio | Sequence number | Mean coverage | Median coverage | Ref. |
|---|---|---|---|---|---|
| DNA-binding | 0.083 | 964 | 0.795 | 0.921 | CLAPE-DB |
| Small molecule-binding | 0.033 | 4888 | 0.870 | 1.000 | CLAPE-SMB |
| Metal ion-binding | 0.015 | 3485 | 0.872 | 1.000 | This work |
| ATP-binding | 0.050 | 390 | 0.838 | 0.917 | E2EATP |
| RNA-binding | 0.186 | 915 | 0.774 | 0.833 | CLAPE |
| Peptide-binding | 0.030 | 3410 | 0.794 | 1.000 | This work |
| PPI homo. | 0.284 | 5649 | 0.709 | 0.727 | DIPS-Plus |
| PPI hetero. | 0.219 | 2591 | 0.734 | 0.778 | DIPS-Plus |
| Enzyme active site | 0.015 | 943 | 0.861 | 1.000 | EasIFA |
| Ion channel | 0.075 | 95 | 0.934 | 1.000 | This work |

104 **Supplementary Table 7 | GO terms used in the ExpGO65 dataset**

| Function label | GO |
|---|---|
| ATP-dependent activity | GO: 0140657 |
| Antioxidant activity | GO: 0016209 |
| Binding | GO: 0005488 |
| Purine ribonucleoside triphosphate binding | GO: 0035639 |
| ATP binding | GO: 0005524 |
| GTP binding | GO: 0005525 |
| Amide binding | GO: 0033218 |
| Antigen binding | GO: 0003823 |
| Carbohydrate binding | GO: 0030246 |
| Carbohydrate derivative binding | GO: 0097367 |
| Chromatin binding | GO: 0003682 |
| Hormone binding | GO: 0042562 |
| Lipid binding | GO: 0008289 |
| Nucleic acid binding | GO: 0003676 |
| DNA binding | GO: 0003677 |
| RNA binding | GO: 0003723 |
| Organic cyclic compound binding | GO: 0097159 |
| Peptide binding | GO: 0042277 |
| Protein binding | GO: 0005515 |
| Protein-containing complex binding | GO: 0044877 |
| Small molecule binding | GO: 0036094 |
| Sulfur compound binding | GO: 1901681 |
| Catalytic activity | GO: 0003824 |
| Cyclase activity | GO: 0009975 |
| Demethylase activity | GO: 0032451 |
| Hydrolase activity | GO: 0016787 |
| Ribonucleoside triphosphate phosphatase activity | GO: 0017111 |
| ATP hydrolysis activity | GO: 0016887 |
| GTPase activity | GO: 0003924 |
| Isomerase activity | GO: 0016853 |

| | |
|---|---|
| Ligase activity | GO: 0016874 |
| Lyase activity | GO: 0016829 |
| Oxidoreductase activity | GO: 0016491 |
| Transferase activity | GO: 0016740 |
| Catalytic activity, acting on a nucleic acid | GO: 0140640 |
| Catalytic activity, acting on a protein | GO: 0140096 |
| Electron transfer activity | GO: 0009055 |
| Molecular adaptor activity | GO: 0060090 |
| Protein-macromolecule adaptor activity | GO: 0030674 |
| Molecular carrier activity | GO: 0140104 |
| Nucleocytoplasmic carrier activity | GO: 0140142 |
| Molecular transducer activity | GO: 0060089 |
| Cytoskeletal motor activity | GO: 0003774 |
| Molecular sequestering activity | GO: 0140313 |
| Protein folding chaperone | GO: 0044183 |
| Receptor activity | GO: 0038024; GO: 0038023 |
| Cargo receptor activity | GO: 0038024 |
| Signaling receptor activity | GO: 0038023 |
| Regulator activity | GO: 0045182; GO: 0140110<br>GO: 0098772; GO: 0140299 |
| Molecular function regulator activity | GO: 0098772; GO: 0140299 |
| ATPase regulator activity | GO: 0060590 |
| Enzyme regulator activity | GO: 0030234 |
| Small molecule sensor activity | GO: 0140299 |
| Signaling receptor regulator activity | GO: 0030545 |
| Transporter regulator activity | GO: 0141108 |
| Transcription regulator activity | GO: 0140110 |
| DNA-binding transcription factor activity | GO: 0003700 |
| Transcription coregulator activity | GO: 0003712 |
| Translation regulator activity | GO: 0045182 |
| Structural molecule activity | GO: 0005198 |

| | |
|---|---|
| Transporter activity | GO: 0005215 |
| Lipid transporter activity | GO: 0005319 |
| Transmembrane transporter activity | GO: 0022857 |
| Carrier activity | GO: 0022857<br>(excluding GO: 0015267) |
| Channel activity | GO: 0015267 |

**Supplementary Table 8 | Prediction performance (functional MCCs) for Word2Function for ExpGO65**

| Function label | Sequence ratio | Sequence number | Functional MCC |
|---|---|---|---|
| ATP-dependent activity | 0.012 | 654 | 0.510 |
| Antioxidant activity | 0.006 | 307 | 0.538 |
| Binding | 0.753 | 42401 | 0.454 |
| Purine ribonucleoside triphosphate binding | 0.015 | 821 | 0.143 |
| ATP binding | 0.010 | 565 | 0.140 |
| GTP binding | 0.005 | 275 | -0.001 |
| Amide binding | 0.006 | 347 | 0.280 |
| Antigen binding | 0.002 | 90 | 0.746 |
| Carbohydrate binding | 0.005 | 292 | 0.183 |
| Carbohydrate derivative binding | 0.027 | 1541 | 0.111 |
| Chromatin binding | 0.015 | 855 | 0.169 |
| Hormone binding | 0.003 | 172 | 0.191 |
| Lipid binding | 0.020 | 1107 | 0.306 |
| Nucleic acid binding | 0.141 | 7916 | 0.537 |
| DNA binding | 0.084 | 4715 | 0.582 |
| RNA binding | 0.059 | 3336 | 0.487 |
| Organic cyclic compound binding | 0.176 | 9903 | 0.451 |
| Peptide binding | 0.005 | 278 | 0.346 |
| Protein binding | 0.638 | 35911 | 0.425 |
| Protein-containing complex binding | 0.046 | 2580 | 0.132 |
| Small molecule binding | 0.082 | 4613 | 0.221 |
| Sulfur compound binding | 0.006 | 349 | 0.144 |
| Catalytic activity | 0.319 | 17973 | 0.722 |
| Cyclase activity | 0.001 | 48 | 0.378 |
| Demethylase activity | 0.001 | 67 | 0.617 |
| Hydrolase activity | 0.098 | 5538 | 0.708 |
| Ribonucleoside triphosphate phosphatase activity | 0.012 | 656 | 0.263 |
| ATP hydrolysis activity | 0.006 | 308 | 0.237 |

| | | | |
|---|---|---|---|
| GTPase activity | 0.006 | 340 | 0.198 |
| Isomerase activity | 0.012 | 654 | 0.690 |
| Ligase activity | 0.012 | 656 | 0.763 |
| Lyase activity | 0.019 | 1072 | 0.656 |
| Oxidoreductase activity | 0.058 | 3250 | 0.751 |
| Transferase activity | 0.127 | 7174 | 0.713 |
| Catalytic activity, acting on a nucleic acid | 0.027 | 1537 | 0.651 |
| Catalytic activity, acting on a protein | 0.088 | 4931 | 0.627 |
| Electron transfer activity | 0.004 | 203 | 0.384 |
| Molecular adaptor activity | 0.026 | 1467 | 0.114 |
| Protein-macromolecule adaptor activity | 0.023 | 1266 | 0.138 |
| Molecular carrier activity | 0.003 | 163 | 0.171 |
| Nucleocytoplasmic carrier activity | 0.001 | 31 | 0.707 |
| Molecular transducer activity | 0.028 | 1569 | 0.691 |
| Cytoskeletal motor activity | 0.001 | 63 | 0.566 |
| Molecular sequestering activity | 0.002 | 136 | 0.297 |
| Protein folding chaperone | 0.002 | 103 | 0.426 |
| Receptor activity | 0.028 | 1577 | 0.663 |
| Cargo receptor activity | 0.001 | 63 | 0.471 |
| Signaling receptor activity | 0.027 | 1523 | 0.674 |
| Regulator activity | 0.098 | 5493 | 0.372 |
| Molecular function regulator activity | 0.047 | 2646 | 0.288 |
| ATPase regulator activity | 0.002 | 95 | 0.603 |
| Enzyme regulator activity | 0.029 | 1637 | 0.253 |
| Small molecule sensor activity | 0.002 | 104 | 0.632 |
| Signaling receptor regulator activity | 0.010 | 579 | 0.421 |
| Transporter regulator activity | 0.003 | 189 | 0.093 |
| Transcription regulator activity | 0.048 | 2699 | 0.461 |
| DNA-binding transcription factor activity | 0.038 | 2158 | 0.498 |
| Transcription coregulator activity | 0.010 | 539 | 0.208 |
| Translation regulator activity | 0.004 | 221 | 0.515 |
| Structural molecule activity | 0.017 | 962 | 0.676 |

| | | | |
|---|---|---|---|
| Transporter activity | 0.061 | 3407 | 0.811 |
| Lipid transporter activity | 0.004 | 207 | 0.343 |
| Transmembrane transporter activity | 0.058 | 3258 | 0.805 |
| Carrier activity | 0.039 | 2174 | 0.812 |
| Channel activity | 0.019 | 1084 | 0.771 |

Sequence ratio refers to the proportion of sequences possessing a particular function relative to the total number of sequences. Sequence number indicates the absolute count of sequences exhibiting this function.

**Supplementary Table 9 | Prediction performance (functional MCCs) for WordTableGO65 for ExpGO65**

| Function label | Sequence ratio | Sequence number | Functional MCC |
|---|---|---|---|
| ATP-dependent activity | 0.012 | 654 | 0.619 |
| Antioxidant activity | 0.006 | 307 | 0.685 |
| Binding | 0.753 | 42401 | 0.457 |
| Purine ribonucleoside triphosphate binding | 0.015 | 821 | 0.307 |
| ATP binding | 0.010 | 565 | 0.267 |
| GTP binding | 0.005 | 275 | 0.344 |
| Amide binding | 0.006 | 347 | 0.371 |
| Antigen binding | 0.002 | 90 | 0.667 |
| Carbohydrate binding | 0.005 | 292 | 0.347 |
| Carbohydrate derivative binding | 0.027 | 1541 | 0.250 |
| Chromatin binding | 0.015 | 855 | 0.292 |
| Hormone binding | 0.003 | 172 | 0.391 |
| Lipid binding | 0.020 | 1107 | 0.384 |
| Nucleic acid binding | 0.141 | 7916 | 0.482 |
| DNA binding | 0.084 | 4715 | 0.537 |
| RNA binding | 0.059 | 3336 | 0.506 |
| Organic cyclic compound binding | 0.176 | 9903 | 0.421 |
| Peptide binding | 0.005 | 278 | 0.377 |
| Protein binding | 0.638 | 35911 | 0.455 |
| Protein-containing complex binding | 0.046 | 2580 | 0.259 |
| Small molecule binding | 0.082 | 4613 | 0.330 |
| Sulfur compound binding | 0.006 | 349 | 0.154 |
| Catalytic activity | 0.319 | 17973 | 0.665 |
| Cyclase activity | 0.001 | 48 | 0.534 |
| Demethylase activity | 0.001 | 67 | 0.771 |
| Hydrolase activity | 0.098 | 5538 | 0.727 |
| Ribonucleoside triphosphate phosphatase activity | 0.012 | 656 | 0.352 |
| ATP hydrolysis activity | 0.006 | 308 | 0.250 |

| | | | |
|---|---|---|---|
| GTPase activity | 0.006 | 340 | 0.432 |
| Isomerase activity | 0.012 | 654 | 0.723 |
| Ligase activity | 0.012 | 656 | 0.802 |
| Lyase activity | 0.019 | 1072 | 0.736 |
| Oxidoreductase activity | 0.058 | 3250 | 0.810 |
| Transferase activity | 0.127 | 7174 | 0.759 |
| Catalytic activity, acting on a nucleic acid | 0.027 | 1537 | 0.748 |
| Catalytic activity, acting on a protein | 0.088 | 4931 | 0.691 |
| Electron transfer activity | 0.004 | 203 | 0.403 |
| Molecular adaptor activity | 0.026 | 1467 | 0.196 |
| Protein-macromolecule adaptor activity | 0.023 | 1266 | 0.206 |
| Molecular carrier activity | 0.003 | 163 | 0.190 |
| Nucleocytoplasmic carrier activity | 0.001 | 31 | 0.353 |
| Molecular transducer activity | 0.028 | 1569 | 0.762 |
| Cytoskeletal motor activity | 0.001 | 63 | 0.640 |
| Molecular sequestering activity | 0.002 | 136 | 0.246 |
| Protein folding chaperone | 0.002 | 103 | 0.210 |
| Receptor activity | 0.028 | 1577 | 0.122 |
| Cargo receptor activity | 0.001 | 63 | 0.769 |
| Signaling receptor activity | 0.027 | 1523 | 0.666 |
| Regulator activity | 0.098 | 5493 | 0.765 |
| Molecular function regulator activity | 0.047 | 2646 | 0.422 |
| ATPase regulator activity | 0.002 | 95 | 0.397 |
| Enzyme regulator activity | 0.029 | 1637 | 0.539 |
| Small molecule sensor activity | 0.002 | 104 | 0.311 |
| Signaling receptor regulator activity | 0.010 | 579 | 0.707 |
| Transporter regulator activity | 0.003 | 189 | 0.581 |
| Transcription regulator activity | 0.048 | 2699 | 0.109 |
| DNA-binding transcription factor activity | 0.038 | 2158 | 0.505 |
| Transcription coregulator activity | 0.010 | 539 | 0.580 |
| Translation regulator activity | 0.004 | 221 | 0.237 |
| Structural molecule activity | 0.017 | 962 | 0.631 |

| | | | |
|---|---|---|---|
| Transporter activity | 0.061 | 3407 | 0.724 |
| Lipid transporter activity | 0.004 | 207 | 0.798 |
| Transmembrane transporter activity | 0.058 | 3258 | 0.515 |
| Carrier activity | 0.039 | 2174 | 0.799 |
| Channel activity | 0.019 | 1084 | 0.811 |

Sequence ratio refers to the proportion of sequences possessing a particular function relative to the total number of sequences. Sequence number indicates the absolute count of sequences exhibiting this function.
